# Contrasting effects of geographic distance, environmental distance, and intraspecific diversity on the performance of a marine invertebrate in common gardens

**DOI:** 10.64898/2026.04.02.716183

**Authors:** Kiran E. Bajaj, Nicole Mongillo, Madeline G. Eppley, Camille A. Rumberger, Zea Segnitz, Shelley Katsuki, Ryan Carnegie, Jessica Small, Katie E. Lotterhos

## Abstract

Restoration and management of natural populations often assume that local genotypes are best suited for transplantation to their local environment. Prioritizing a single local genotype, however, contrasts with the framework of maximizing intraspecific diversity to increase population resilience to environmental change. Local populations may also become maladapted to a rapidly changing environment, motivating alternative frameworks that instead minimize environmental distance between source and transplantation sites. Here, we tested the predictive power of the *local is best*, *maximize intraspecific diversity*, and *minimize environmental distance* frameworks on the survival and growth of Eastern oyster (*Crassostrea virginica*) genotypes in field common gardens that differed in salinity and disease pressure. Although a genome scan revealed patterns of adaptation to disease, heat stress, and salinity among source populations, we did not find strong support for the *local is best* framework: geographically distant southern genotypes performed comparably to local selection lines and a local wild population. Higher genetic diversity within monocultures was associated with higher survival, yet highly diverse polycultures survived at lower rates than the best-performing monocultures, providing mixed support for the *maximize intraspecific diversity* framework. Temperature and salinity of the environments-of-origin of parents predicted the survival of their offspring in common gardens, but the relationship between survival and environmental distance was context-dependent, leading to mixed support for the *minimize environmental distance* framework. Together, these results demonstrate that no single framework reliably predicted transplantation success, suggesting that effective management strategies may need to integrate genomic and environmental lines of evidence to guide genotype selection.

## Introduction

Ecologists and evolutionary biologists have traditionally emphasized different mechanisms underlying the dynamics of foundation species. Ecologists have examined how intraspecific trait diversity—arising through niche partitioning or selective survival—shapes ecosystem function (Bolnick et al., 2011; Des Roches et al., 2018; Govaert et al., 2024; Loreau et al., 2001; Moran et al., 2016; Palacio et al., 2025; Raffard et al., 2019). Under niche partitioning, genotypes occupy distinct ecological roles that limit competition and promote productivity (Svanbäck & Bolnick, 2006). Under selective survival, genetic diversity increases the chance that disturbance-tolerant genotypes will sustain productivity (Fargione et al., 2007). In contrast, evolutionary biologists have tested how genotypes perform across environments and have identified genomic regions associated with selection and adaptive traits (Barrett & Hoekstra, 2011; Bernatchez et al., 2024; Hoban et al., 2016; Lasky et al., 2023; Rellstab et al., 2015; Savolainen et al., 2013), an approach that maximizes productivity by matching genotypes to their adaptive environment. Both fields share the goal of identifying guiding principles for selecting or combining genotypes to maximize productivity.

Many applications prioritize local genotypes as source material, a guiding principle we refer to as the *local is best* framework. Local adaptation, which occurs when populations have higher fitness in sympatry than in allopatry (Blanquart et al., 2013; Kawecki & Ebert, 2004), underlies this framework. In practice, forestry often restricts planting stock to local sources (Boshier et al., 2015; Nolan et al., 2023), and conservation efforts prioritize local or regional gene pools (Bucharova et al., 2019; Jones, 2013). Local sourcing remains less common in aquaculture due to assumed high gene flow in marine species (Conover, 1998), though evidence for local adaptation is accumulating (DuBois et al., 2022; Sanford & Kelly, 2011). Importantly, the definition of “local” varies with how practitioners use environmental, genetic, demographic, and geographic data (Boshier et al., 2015).

Rapid climate change is causing populations to become maladapted to their local environments (Gougherty et al., 2021; Jeffery et al., 2024; Parmesan, 2006; Sgrò et al., 2011; Tigano et al., 2024), challenging the *local is best* framework and motivating a shift towards climate matching. Under a strategy we term the *minimize environmental distance* framework, donor populations are chosen based on environmental similarity between source and recipient sites (Jones, 2013; Khazaei et al., 2013). This approach underlies assisted gene flow, in which pre-adapted individuals are relocated to enhance adaptation in recipient populations through genetic admixture (Sgrò et al., 2011; Aitken & Whitlock, 2013; Aitken & Bemmels, 2016). Recent studies report no evidence of outbreeding depression from assisted gene flow (reviewed in Ralls et al. 2020, Kelly et al. 2021, Pregler et al. 2023), yet concerns persist about using genetically distant sources even when environments align (Byrne et al., 2011; Grummer et al., 2022; Weeks et al., 2011).

Ecological and evolutionary perspectives also recognize the benefits of intraspecific diversity, either for supporting resistance to disturbance, productivity, and community processes (Crutsinger et al., 2006; Govaert et al., 2024; Hughes & Stachowicz, 2004; Raffard et al., 2019; Reusch et al., 2005), or as the raw material for evolutionary change and long-term persistence under environmental stress (Barrett & Schluter, 2008; Bay et al., 2018; Waldvogel et al., 2020). These perspectives converge on a *maximize intraspecific diversity* framework, often applied in restoration (Kettenring et al., 2014; Stockwell et al., 2016). However, methods for generating diversity have not been systematically compared. For example, diversity can be generated by mixing individuals from different populations or by producing novel crosses which may benefit from hybrid vigor (Stelkens et al., 2014) or suffer from outbreeding depression (Edmands, 1999).

Genotype selection for agriculture, aquaculture, and restoration is complicated by these contrasting ecological and evolutionary frameworks that have not been compared in a single study. Here, we address this gap using common garden field experiments to assess the productivity of a foundational reef-building species, the Eastern oyster (*Crassostrea virginica*), in aquaculture and field settings. Through a collaboration with an aquaculture breeding center, we had the unique opportunity to raise offspring from both local and geographically/genetically distant groups at two common garden sites differing in salinity and disease pressures. We tracked proxies of fitness from larval development through field grow-out and integrated population genomic data to identify genomic regions underlying differential survival. This design allowed us to directly test the *local is best*, *minimize environmental distance*, and *maximize intraspecific diversity* frameworks (Table 1).

**Table 1.**
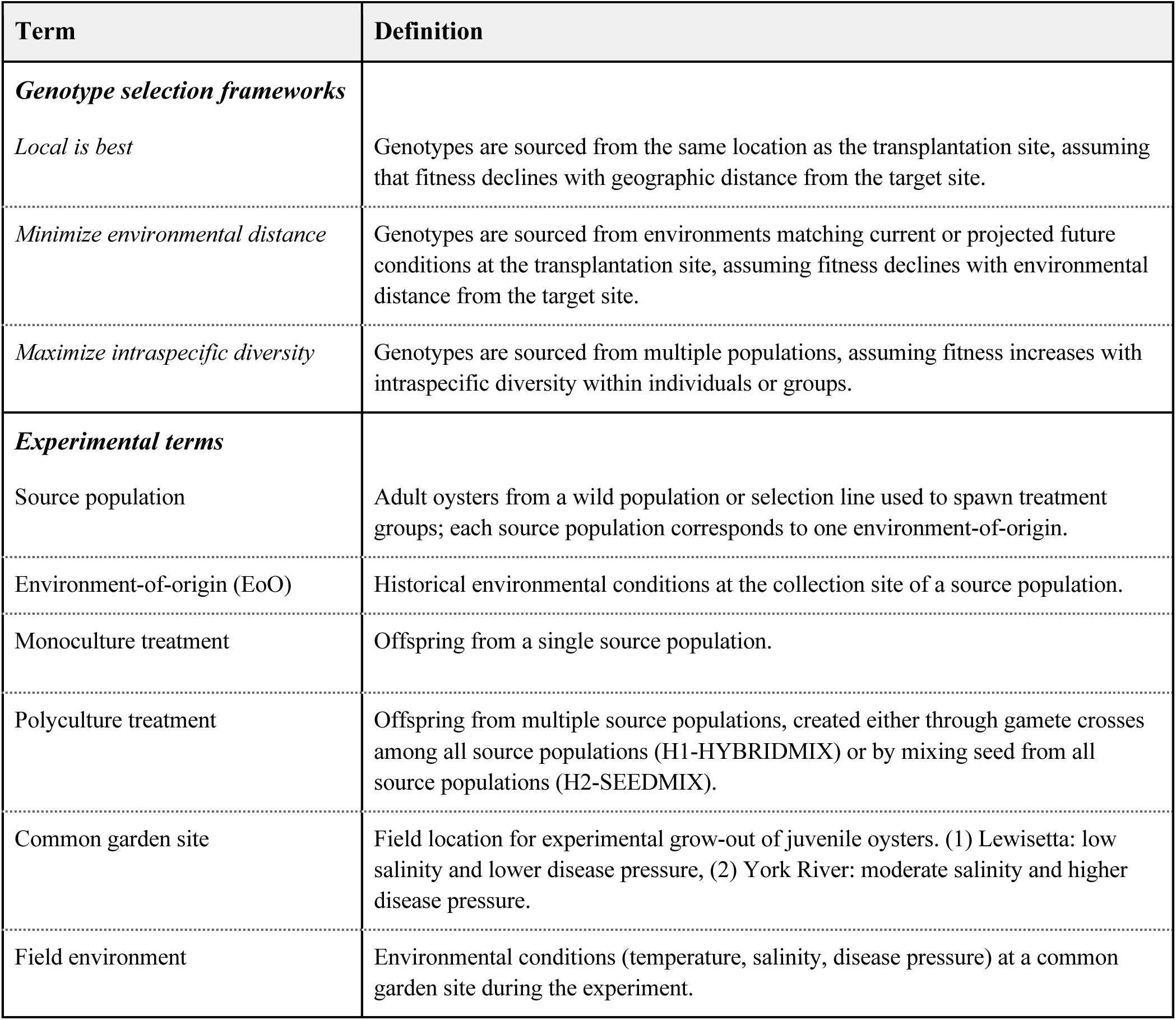
Terms and definitions. Descriptions of genotype selection frameworks, experimental treatments, and sites.

## Methods

### Study system

Eastern oysters adapt to salinity, temperature, and disease across their extensive range from Texas, USA to Nova Scotia, Canada (Eppley et al., 2026; Fuqua & Brooke, 2025; Griffiths et al., 2021; Shumway, 1996). The protozoan pathogens MSX (*Haplosporidium nelsoni*) and Dermo (*Perkinsus marinus*) are major sources of mortality (Ragone Calvo et al., 2003). Some populations or selection lines have evolved resistance or tolerance to at least one of these pathogens, but mortality remains high under heavy infections (Casas et al., 2017; Proestou et al., 2019; Jiang et al., 2024; Wang et al., 2025, see also “Methods: Environmental data”). Disease prevalence and intensity generally increase with salinity and temperature (Ewart & Ford, 1993; Ford & Haskin, 1982; Soniat et al., 2006).

### Experimental treatments

Our experimental design consisted of ten treatments (eight monocultures and two polycultures, Table 2) grown at two sites in the Chesapeake Bay, Virginia, USA. The York River site is characterized by moderate salinity and high disease pressure; the Coan River site (“Lewisetta”) is characterized by relatively lower salinity and lower disease pressure (Figure 1A, 1C).

**Figure 1.**
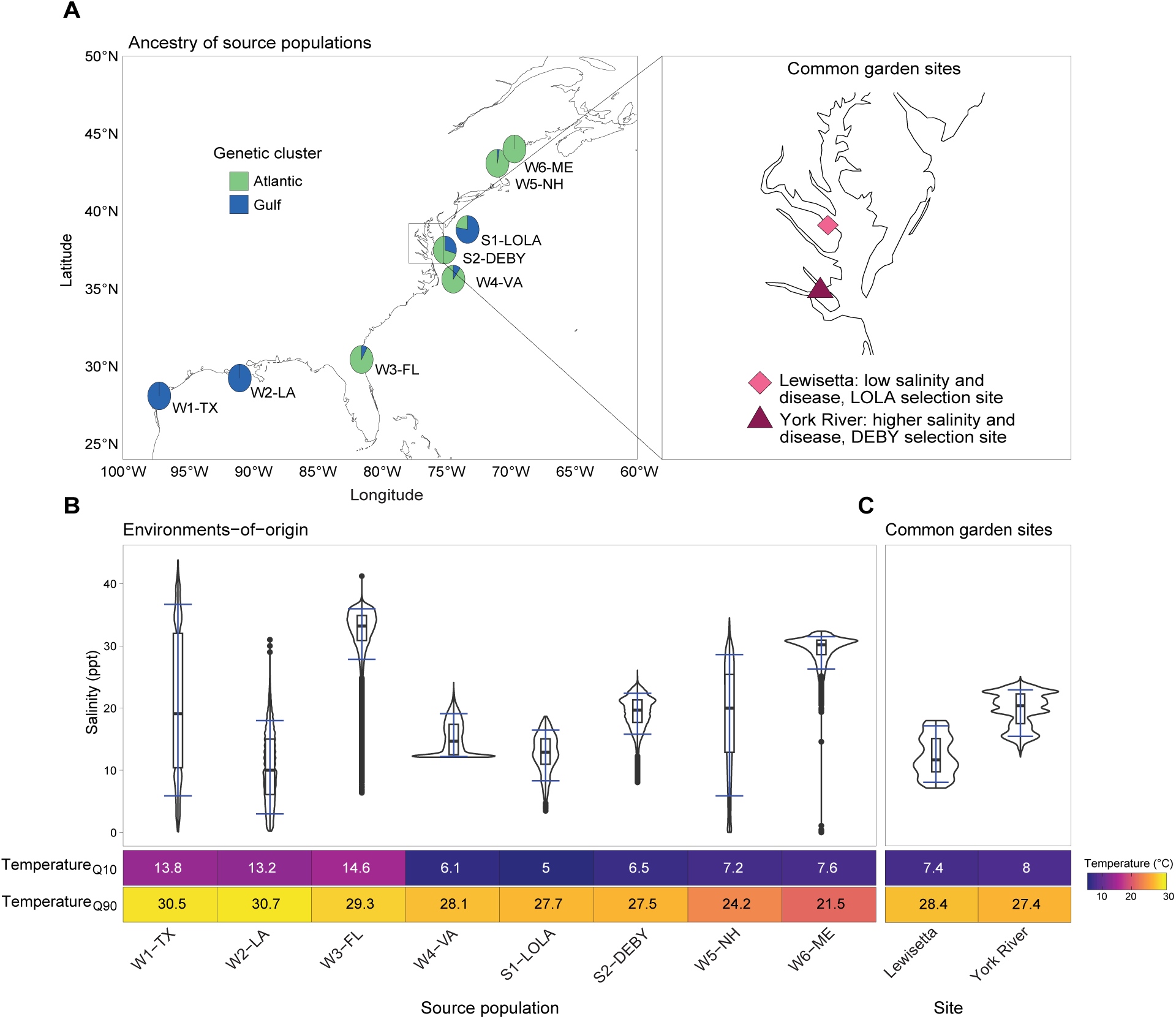
Source populations. (A) Collection locations of each source population. Pie charts indicate the proportion of Gulf (blue) and Atlantic (green) ancestry based on sNMF analysis with support for K = 2 genetic clusters. Inset: common garden sites in the Chesapeake Bay: Lewisetta (diamond) and the York River (triangle). (B) Environmental conditions at source populations’ environments-of-origin. Violin plots of raw salinity (black) and 0.1 and 0.9 quantiles (Salinity_Q10_ and Salinity_Q90_; blue bars), with heatmaps of 0.1 and 0.9 temperature quantiles (Temperature_Q10_ and Temperature_Q90_) below. (C) Salinity violin plot and temperature heatmaps for common garden sites.

**Table 2.** Experimental treatments. Ten treatments representing wild monocultures (W1-W6), selection line monocultures (S1-S2), and polycultures (H1-H2). Relative geographic origin indicates position relative to common garden sites in the Chesapeake Bay. Primary genetic ancestry reflects Gulf or Atlantic lineage origin. Polyculture treatments combined genetic material (H1-HYBRIDMIX) or nursery seed (H2-SEEDMIX) from all monoculture source populations in equal proportions; detailed crossing and mixing protocols are described in Methods.

| Treatment | Treatment type | Source population | Geographic origin | Relative geographic origin | Primary genetic ancestry | Latitude | Longitude |
| --- | --- | --- | --- | --- | --- | --- | --- |
| W1-TX | Wild monoculture | W1-TX | Copano Bay, Texas | Southern | Gulf | 28.096000 | -97.174000 |
| W2-LA | Wild monoculture | W2-LA | Sister Lake, Louisiana | Southern | Gulf | 29.239925 | -90.911362 |
| W3-FL | Wild monoculture | W3-FL | Timucuan Preserve, Florida | Southern | Atlantic | 30.440030 | -81.436380 |
| W4-VA | Wild monoculture | W4-VA | James River, Virginia | **Local (to both gardens)** | Atlantic | 37.1501163 | -76.638923 |
| S1-LOLA | Selection line monoculture | S1-LOLA | Coan River, Virginia | **Local (to Lewisetta)** | Gulf | 37.98030 | -76.46190 |
| S2-DEBY | Selection line monoculture | S2-DEBY | York River, Virginia | **Local (to York River)** | Atlantic | 37.249107 | -76.497032 |
| W5-NH | Wild monoculture | W5-NH | Great Bay, New Hampshire | Northern | Atlantic | 43.053746 | -70.911544 |
| W6-ME | Wild monoculture | W6-ME | Hog Island, Maine | Northern | Atlantic | 44.01330 | -69.541689 |
| H1-HYBRIDMIX | Polyculture with outcrossing | All source populations | Created by crossing sperm from each individual male from with a pool of eggs in equal proportions from all females from all the monoculture treatments. The resulting zygotes were then pooled and raised in the hatchery. |  |  |  |  |
| H2-SEEDMIX | Polyculture | All source populations | Prior to field deployment of experimental treatments, created by mixing nursery seed from each monoculture treatment in equal proportions. |  |  |  |  |

Source populations for monoculture treatments included six wild populations and two proprietary broodstock lines (DEBY and LOLA) from the Aquaculture Genetics and Breeding Technology Center (ABC), Virginia Institute of Marine Science (VIMS). Wild populations spanned the species’ native range, which is structured into distinct genetic clusters separated by the Florida peninsula (Figure 1A; Figure S1; Reeb & Avise, 1990, Puritz et al., 2022).

Populations included oysters from: a variable salinity site in Texas (W1-TX), a low salinity site in Louisiana (W2-LA), a high salinity site on the east coast of Florida (W3-FL), a moderate salinity site in the James River, Virginia (W4-VA), a variable salinity site in New Hampshire (W5-NH), and a high salinity site in Maine (W6-ME) (Figure 1B; Table 2). W4-VA was considered local to both common garden sites due to its geographic proximity and intermediate salinity (Figure 1B, 1C; Table 2).

The selection line DEBY originated from crosses between wild Delaware Bay and Virginia oysters in 1987, with some sub-lines later crossed with Louisiana oysters and consolidated into a “superline” in 2008-2009. DEBY has since undergone 8 generations of selection in the York River (Puritz et al., 2022; Ragone Calvo et al., 2003). LOLA was established in 2008 by crossing Louisiana oysters from multiple sites (80%) with DEBY (20%) and has been selected for at least 4 generations at Lewisetta (Puritz et al. 2022). In this study, DEBY is considered local to the York River and LOLA to Lewisetta.

Broodstock were collected September-November 2022 and conditioned at the Acuff Center for Aquaculture at VIMS until spawning on May 9, 2023 (Supp. Methods: broodstock conditioning). To minimize sperm competition (Levitan, 2004), oysters were strip-spawned and sperm from 10 males per monoculture treatment was individually crossed with pooled eggs from 10 equally represented females. Zygotes were then combined within each monoculture. The first polyculture treatment (H1-HYBRIDMIX) was created by crossing sperm from each experimental male with pooled eggs from all experimental females, generating novel genetic combinations among source populations (Supp. Methods: spawning and fertilization). Larvae were reared following standard hatchery (days 0-30) and nursery (days 30-77) procedures (Supp. Methods: larval rearing). On day 77, the second polyculture treatment (H2-SEEDMIX) was created by mixing equal proportions of seed from the eight monocultures, increasing genetic diversity while preserving original genetic compositions of each source population.

### Field deployment and monitoring

Seed from each treatment were split into six bags (1,200 per bag). Three replicate bags were deployed in bottom cages at each common garden site in August 2023 and assessed in November 2023, May 2024, and November 2024. At the first assessment, 40 oysters per bag were individually tagged. At each assessment, live and dead oysters were counted and shell length (longest distance between hinge and bill) and width (longest distance perpendicular to length) were measured for tagged individuals. Five oysters per bag were frozen for condition index analysis (tissue dry weight / shell dry weight ×100; Rainer and Mann 1992). To reduce density-dependent effects, bags were thinned to 225 oysters in May 2024 (Marshall & Dunham, 2013).

Cumulative survival was calculated at each timepoint (Supp Methods: survival measurements). Response variables included treatment-level survival (3 bags per treatment per site), individual size (*n* = 40 tagged oysters per bag), and individual condition index (*n* = 5 oysters per bag per timepoint).

### Environmental data

*Source population environment-of-origin:* For each source population, hourly (or finer) temperature and salinity data spanning > 8 years were obtained from a nearby buoy. At sites without continuous onsite data, data was obtained from a more distant buoy and adjusted using limited onsite data (Table S1; Supp. Methods: environmental data). We characterized each environment-of-origin by the 0.1 and 0.9 quantiles of temperature and salinity (Temperature_Q10_, Temperature_Q90_, Salinity_Q10_, Salinity_Q90_, Table S2), which captured extreme conditions likely to impose selection and were not significantly correlated (pairwise r < 0.7, Figure S2).

*Historical disease exposure:* Although we didn’t quantify disease at the environments-of-origin, we can infer historical disease exposure based on geographic location and salinity. Dermo occurs throughout the species range, present at least as far north as northern Maine (Boeck, 2018; Marquis et al., 2020), whereas MSX is common along the Atlantic coast (Ewart & Ford, 1993; Marquis et al., 2015) but is reported as absent (Boeck et al., 2019) or rare (Ulrich et al., 2007) west of the Florida peninsula (hereafter: “Gulf”). Prevalence and intensity of both diseases generally increases with salinity and temperature (Burreson and Andrews 1988, Silvy et al. 2020, Gignoux-Wolfsohn et al. 2021), but these relationships vary regionally within the Chesapeake Bay (Kachmar et al., 2025). Dermo generally causes more mortality than MSX (Carnegie & Burreson, 2011). We therefore expect W1-TX, W3-FL, W4-VA, and S2-DEBY to have experienced the most frequent Dermo exposure, and W3-FL, W4-VA, and S2-DEBY (followed by W5-NH and W6-ME) to have experienced the most frequent MSX exposure.

*Experimental environment:* Daily salinity and temperature at each common garden site were obtained from nearby buoys throughout the experiment (Table S1). Naive sentinel oysters were deployed between May and December at each common garden site and collected every 2 weeks to quantify disease pressure (n = 25/site/date). Mantle tissue samples from sentinel oysters were cultured with Ray’s fluid thioglycollate medium (RFTM) and stained with Lugol’s iodine solution for microscopic evaluation of individual-level infection intensity and population-level disease prevalence (Audemard et al., 2008; Ray, 1952).

### Genetic analysis

Gill tissue was collected from each parent (*n* = 160) after spawning and stored in 95% ethanol at-80°C. DNA was extracted using the Qiagen DNeasy Blood and Tissue Kit (Supp. Methods: DNA extraction) and shipped to Neogen Genomics (Lincoln, NE) for genotyping on a 200K ThermoFisher Affymetrix Axiom SNP array derived from a 600K array (Gómez-Chiarri et al., 2015; Guo et al., 2023; Modak et al., 2021; Puritz et al., 2024) aligned to the haplotig-masked *Crassostrea virginica* reference genome (C_virginica-3.0, GCA_002022765.4; Supp. File 1).

Raw genotypes were processed and filtered in R v4.4.2 (R Core Team 2025; Supp. Methods: genotype filtering). Missing SNPs were imputed using LEA v3.6.0 (Frichot & François, 2015) with K = 2 ancestral groups to produce a *full SNP set*. We thinned SNPs for linkage disequilibrium (LD) for population structure analysis and neutral parameterization of genome scans (Lotterhos, 2019) using bigsnpr v1.11.3 (Privé et al., 2018). Ancestry was inferred on the *thinned SNP set* using sNMF(). We tested K = 1-20 and selected the value with the lowest cross-entropy across 10 repetitions.

## Statistical analysis

We used statistical analyses to answer specific questions about the survival, growth, and condition index of monoculture and polyculture treatments (summarized in Table 3). Model assumptions were assessed and variables were transformed as needed (Supp. Methods: statistical analysis). For models with multiple explanatory variables, we used Akaike’s Information Criterion (AIC) to select the simplest and best-fitting model.

**Table 3.**
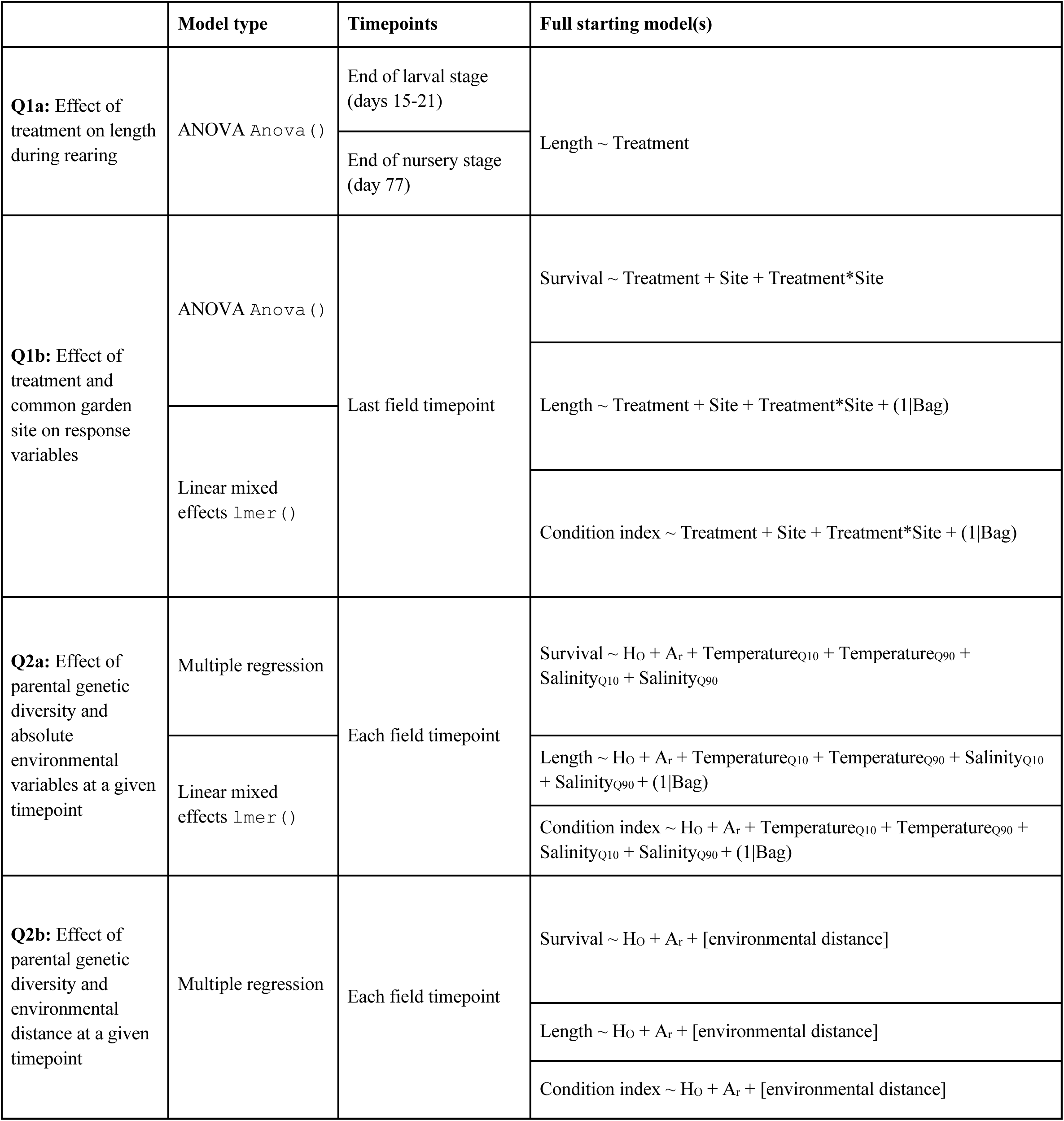
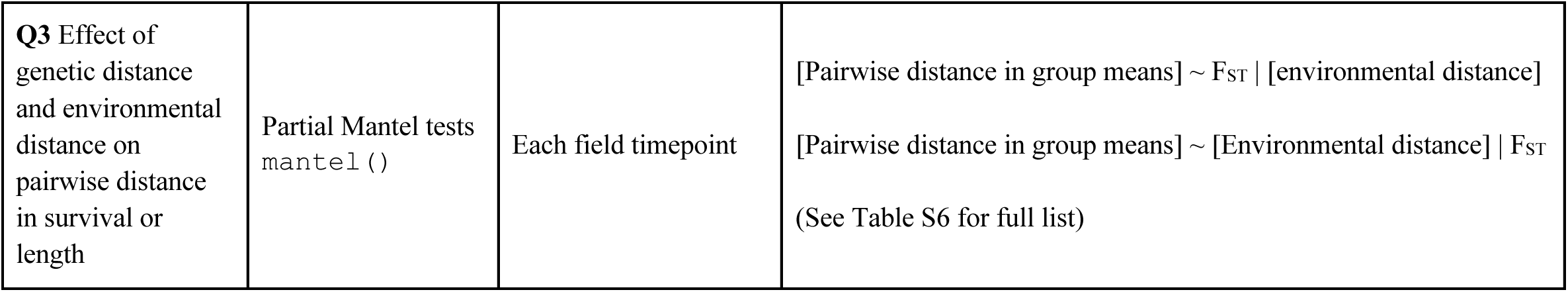
Overview of the statistical models used in this study for each study question.

### Q1: How do survival and growth vary among treatments in the hatchery and at different field sites?

We tested for differences in shell length among treatments at the end of the larval stage (days 15-21) and nursery stage (day 77) using ANOVA (Table 3, Q1a). At the final field timepoint (November 2024), we modeled bag-level survival as a function of treatment, common garden site, and their interaction using 2-way ANOVA (Table 3, Q1b). We modeled shell length and condition index as a function of treatment, common garden site, and their interaction with bag as a random effect using lme4 v1.1.37 (Bates et al., 2015) (Table 3, Q1b). Pairwise differences were tested with Tukey-Kramer post hoc tests.

### Q2: To what extent does parental genetic diversity and environment-of-origin predict offspring performance?

For the monoculture treatments, we took two regression approaches. First, to test if the response variables were related to genetic diversity and the temperature and salinity at the source populations’ environment-of-origin, we modeled bag-level survival (field only), shell length (hatchery, nursery, and field), or condition (field only) as a function of heterozygosity, allelic richness, Temperature_Q10_, Temperature_Q90_, Salinity_Q10_, and Salinity_Q90_ (Table 3, Q2a). Second, we replaced absolute environmental conditions with pairwise Euclidean environmental distance between each source populations’ environment-of-origin and the common garden site, calculated on standardized temperature and salinity quantiles (Table 3, Q2b). Heterozygosity and allelic richness of each source population were calculated on the *full SNP set* with hierfstat v.0.5.11 (Goudet, 2005) using a rarefaction level of 40, based on the population size of 20 individuals multiplied by two to account for diploidy.

### Q3: To what extent do pairwise genetic and environmental differences predict pairwise performance differences in growth or survival?

We used partial Mantel tests with Spearman’s correlation and 10,000 permutations (vegan v2.7.1; Oksanen et al., 2025) to test whether pairwise differences in shell length or survival were predicted by genetic distance (pairwise *F_ST_*, Weir & Cockerham, 1984, calculated on the *thinned SNP set* with hierfstat) while controlling for environmental distance or by environmental distance while controlling for genetic distance (Table 3, Q3).

### Q4: What genomic regions are putatively under spatially heterogeneous selection?

To explore the genetic basis of treatment-level differences observed in the field, we conducted a genome scan using OutFLANK v0.2 (Whitlock & Lotterhos, 2015). We calculated per-locus values of *FST* among source populations with neutral parameters estimated from the *thinned SNP set* (trim fractions = 0.05; minimum heterozygosity = 0.1). Outliers were identified in the *full SNP set* using a *q*-value < 0.01. SNPs were annotated by intersecting genomic coordinates with gene models from the *C. virginica* genome v3.0 GFF3 (GCA_002022765.4, release 62) using GenomicRanges v1.58.0 (Lawrence et al., 2013) and biomaRt v2.62.1 (Durinck et al., 2009). Gene ontology enrichment was conducted with ShinyGO 0.82 (Ge et al., 2020).

## Results

### Environmental data

Salinity_Q10_ and Salinity_Q90_ spanned high (25-36 ppt; W3-FL, W6-ME), moderate (10-25 ppt; W4-VA, S2-DEBY), low (3-18 ppt; W2-LA, S1-LOLA), and highly variable (5-37 ppt; W1-TX, W5-NH) salinity conditions. (Figure 1B). Only Gulf sites experienced temperatures exceeding 30°C. High-temperature exposure (Temperature_Q90_) declined with latitude, from 30-31°C in the south to 21-25°C in the north (Figure 1B). Low-temperature exposure (Temperature_Q10_) was similar between Chesapeake Bay and northern Atlantic sites (6-8°C). Southern sites experienced milder lows (13-15°C). Based on established relationships between temperature, salinity, and disease pressure (Burreson and Andrews 1988, Silvy et al. 2020, Gignoux-Wolfsohn et al. 2021), W1-TX, W3-FL, W4-VA, and S2-DEBY were assumed to have experienced the highest historical disease pressure due to consistent salinity above 15 ppt and temperature above 27°C (Soniat et al., 2006).

The Lewisetta and York River common garden sites experienced similar temperatures but significantly different salinities (8.7-16.6 ppt vs. 15.4-22.9 ppt, respectively; Figure 1C). Dermo pressure was lower at Lewisetta (5-64% prevalence) and maximal at the York River, reaching 100% (Figure 2A). In 2023, when oysters were approximately six months old, an MSX outbreak at the York River site reached 53% prevalence (Figure 2B, Tarnowski 2024).

**Figure 2.**
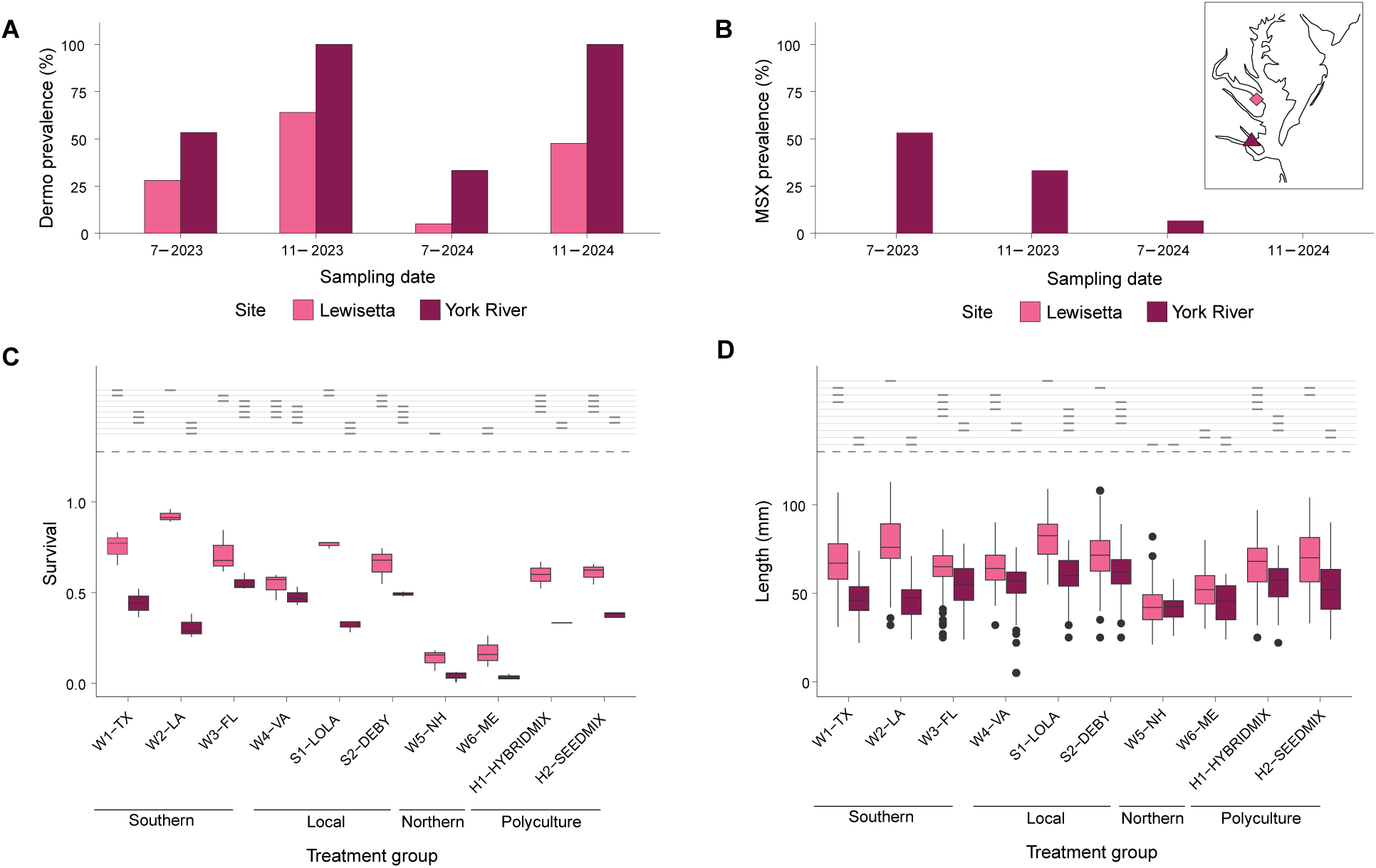
Field performance. (A) Dermo prevalence (%) quantified by Ray’s fluid thioglycollate medium and (B) MSX prevalence (%) quantified by histopathology at the Lewisetta and York River common garden sites during the field experiment. MSX was not detected at Lewisetta. Boxplots of (C) survival and (D) shell length of each treatment group at the final field monitoring event (November 2024) at the Lewisetta (lighter boxes) and York River (dark boxes) common garden sites. Horizontal lines above the boxes indicate significant pairwise differences: groups that share a line at the same height do not have a statistically significant difference.

### Genetic analysis and population structure

After filtering for >10% missing data across all loci (n = 6,929) and individuals (n = 0), 188,096 SNPs remained. Filtering for minor allele frequency > 0.05 yielded 154,102 SNPs across 160 parents (*full SNP set*; Supp. File 1). LD pruning reduced this to 105,356 SNPs (*thinned SNP set*; Supp. File 1). sNMF ancestry analysis supported two genetic clusters (K = 2) separating Gulf and Atlantic source populations (Figure 1A; Figure S1). Pairwise *F_ST_* was low within regions (<0.001 within the Gulf; ∼0.02 within the Atlantic) and reached a maximum of 0.05 between the western Gulf and northern Atlantic.

### Southern genotypes outperformed northern genotypes in the field

Mono/polyculture treatment significantly affected shell length at the end of both the larval stage (days 15-21; ANOVA, df = 8, F = 28.14, p < 0.001) and the nursery stage (day 77; ANOVA, df = 8, F = 51.03, p < 0.001). W6-ME oysters were the largest during the larval stage but among the smallest by the end of the nursery stage, while the low salinity selection line (S1-LOLA) had intermediate larval shell length but was significantly larger than all other treatments by the end of the nursery stage. W5-NH oysters were consistently the smallest (Supp. Figure S3).

At the final field timepoint, common garden site, treatment, and their interaction significantly influenced survival and shell length, consistent with GxE effects (Tables S3, S4). Common garden site and treatment also affected condition (Table S5). Performance was consistently poorer at the York River site than at Lewisetta across all metrics (Figure 2C, 2D; Figure S4).

Northern Atlantic monocultures (W5-NH, W6-ME) performed poorly at both sites, with the lowest survival (mean = 3.44-17.11%, Figure 2C) and smallest shell lengths (mean = 41.74-52.34 mm, Figure 2D). At Lewisetta, Gulf and southern-origin monocultures (W1-TX, W2-LA, S1-LOLA, W3-FL) had the highest survival (mean = 71.28-92.21%, Figure 2C) and largest sizes (mean = 63.18-81.18 mm, Figure 2D). In the York River, southern monocultures (W1-TX, W3-FL, W4-VA, and S2-DEBY) had the highest survival (mean = 44.20-55.45%, Figure 2C), consistent with their assumed history of higher Dermo exposure. The local selection line (S2-DEBY) had the largest shell lengths in the York River (mean = 61.55 mm, Figure 2D).

The two polyculture treatments (H1-HYBRIDMIX, H2-SEEDMIX) did not differ from each other in any metric and consistently performed worse than the best monocultures at both common garden sites (Figure 2C, 2D; Figure S4).

### Source population genetic diversity and environment-of-origin predicted offspring survival

We first tested whether performance of monocultures was predicted by genetic diversity and absolute conditions at the source populations’ environments-of-origin (Table 3, Q2a).

Temperature_Q90_ was strongly correlated with heterozygosity (*r* = 0.72), and allelic richness (*r* = 0.74) (Supp. Figure S2) and was excluded from models to avoid multicollinearity, but is displayed alongside heterozygosity and allelic richness in Figure 3. Survival and shell length were positively associated with source population heterozygosity, allelic richness (and higher Temperature_Q90_), and higher minimum salinity (Salinity_Q10_) (Figure 3A, 3C; Supp. Figures S5, S6). These relationships were generally consistent across timepoints and common garden sites, with few exceptions (Supp. File 3). Predictors of condition were weak or inconsistent (Supp. Figure S7; note that we did not observe spawning activity at sampling timepoints).

**Figure 3.**
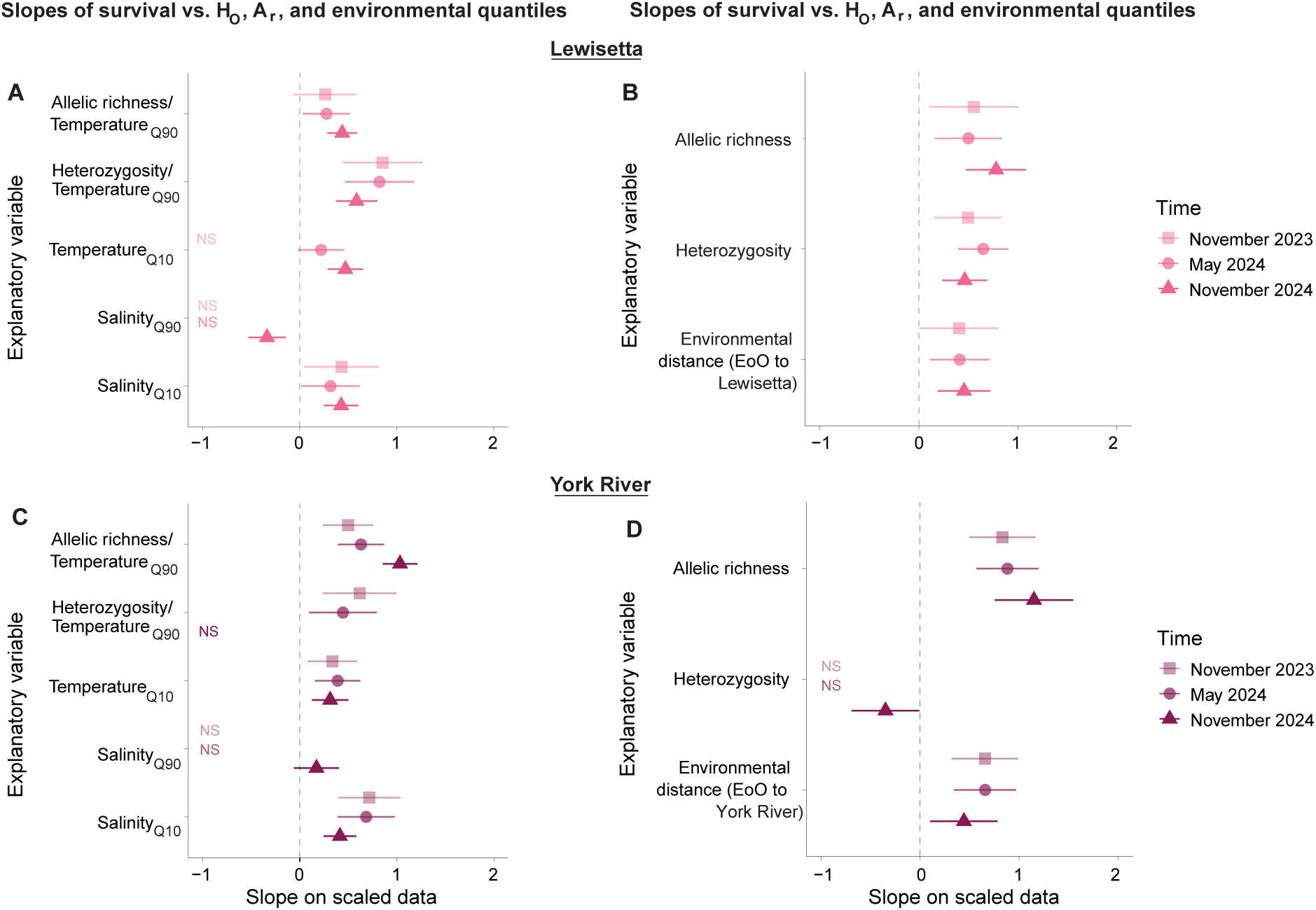
Relationships among field survival, source population genetic diversity, and source environment. Slope ± 2SE of scaled survival at Lewisetta at three field monitoring events vs. heterozygosity, allelic richness, and either (A) absolute conditions at source populations’ environments-of-origin (EoO) or (B) environmental distance from EoOs to the Lewisetta common garden site. Slope ± 2SE of scaled survival in the York River at three field monitoring events vs. heterozygosity, allelic richness, and either (C) absolute conditions at source populations’ EoO or (D) environmental distance from EoOs to the York River common garden site.

We also tested whether environmental distance between common garden sites and source populations’ environments-of-origin predicted performance in the field. Consistent with the previous analysis, allelic richness was positively associated with survival at all timepoints at both sites (Figure 3B, 3D). At Lewisetta, heterozygosity was positively related to survival through time, whereas this relationship was absent or marginally negative at the York River (Figure 3B, 3D). The correlation between survival and environmental distance was negative at both common garden sites, consistent with the *minimize environmental distance* framework (Supp. Figure S8). However, in a full model including genetic diversity metrics, the relationship between survival and environmental distance was positive through time at both common garden sites, which is unexpected under the *minimize environmental distance* framework (Figure 3B, 3D). This latter pattern was largely driven by W3-FL, which had a high environmental distance (due to the much higher salinity at the EoO) but relatively higher survival after accounting for genetic diversity (Supp. Figure S9). Relationships between predictors and shell length or condition were variable through time and across sites, with no consistent pattern (Supp. Figure S5, S7, S8, S9).

### Pairwise genetic and environmental distances did not predict pairwise differences in performance

After correcting for multiple comparisons, no partial Mantel test showed a significant relationship between pairwise differences in survival or shell length and either genetic or environmental distance (Figure 4, Figure S10, Table S6).

**Figure 4.**
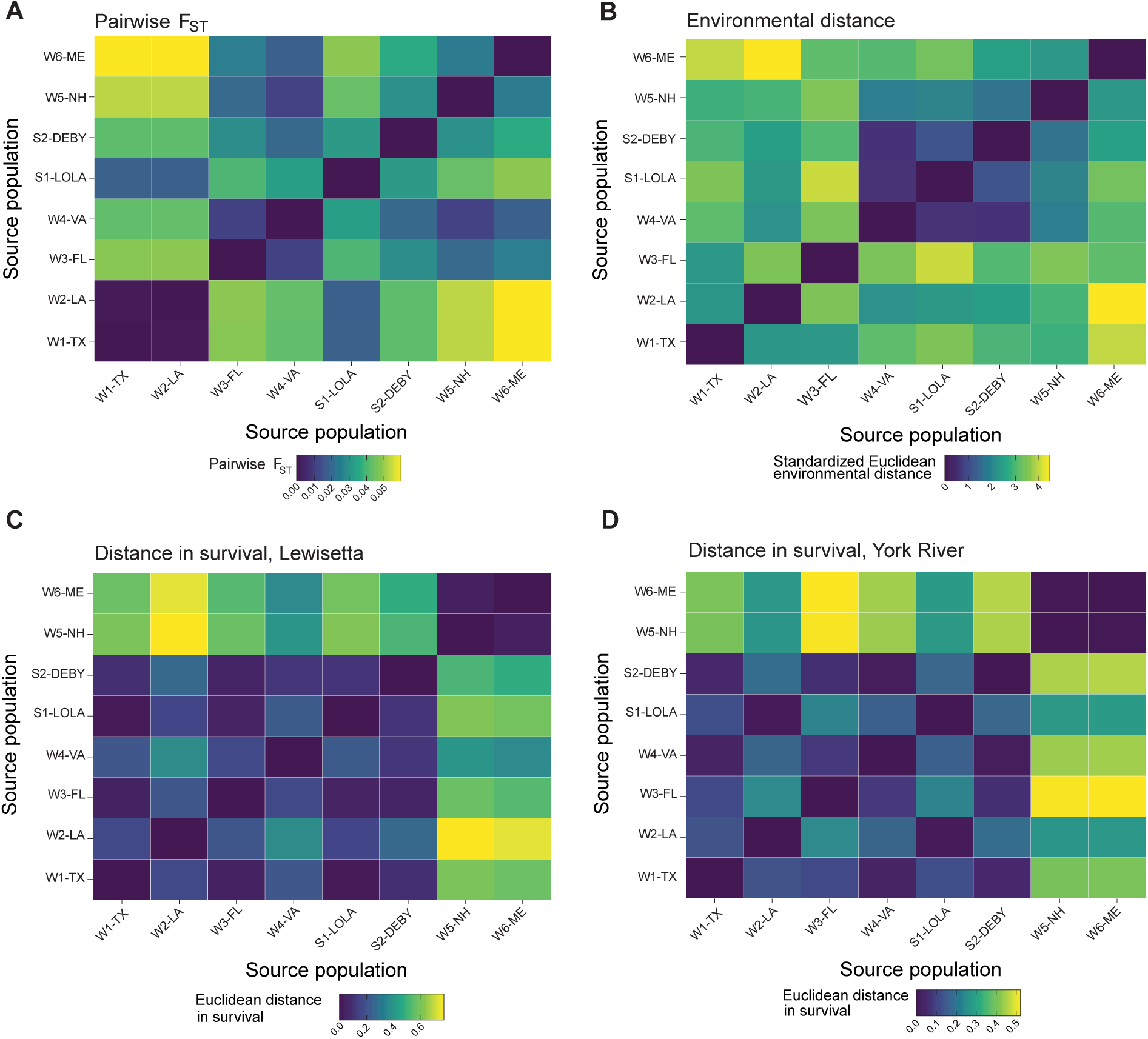
Pairwise distances. (A) Heatmap of pairwise *F_ST_* calculated on SNPs thinned for LD and (B) pairwise Euclidean distance between environments-of-origin based on standardized 0.1 and 0.9 quantiles of temperature and salinity. Euclidean distance in survival between treatment groups at the final field timepoint at (C) Lewisetta and (D) the York River. Darker colors indicate small pairwise differences and lighter colors indicate greater pairwise differences.

### Genomic outliers implicated disease resistance, stress response, and growth pathways

OutFLANK identified 6,928 outlier SNPs, of which 5,041 were successfully annotated, representing 1,634 unique genes (Figure 5A, Supp. File 5). Several outliers mapped to genes previously linked to Dermo resistance, including LOC111118916, LOC111122703, LOC111122700, and LOC111123901 (Wang et al. 2025; Supp. File 4). The strongest outlier (*F_ST_* = 0.98) mapped to LOC111124621, annotated as myosin heavy chain (striated muscle-like) and enriched for ATP binding (GO:0005524). Another myosin heavy chain gene (LOC111118916, non-muscle isoform), previously associated with Dermo resistance (Wang et al. 2025), also showed high differentiation (*F_ST_* = 0.69). Outliers were also detected in advillin (LOC111122703) and a nearby Dnaj homolog in the HSP40 heat shock family (LOC111122700), but neither were assigned GO terms. A tumor suppressor gene (LOC111123901) was also an outlier, associated with signal transduction (GO:0007165).

**Figure 5.**
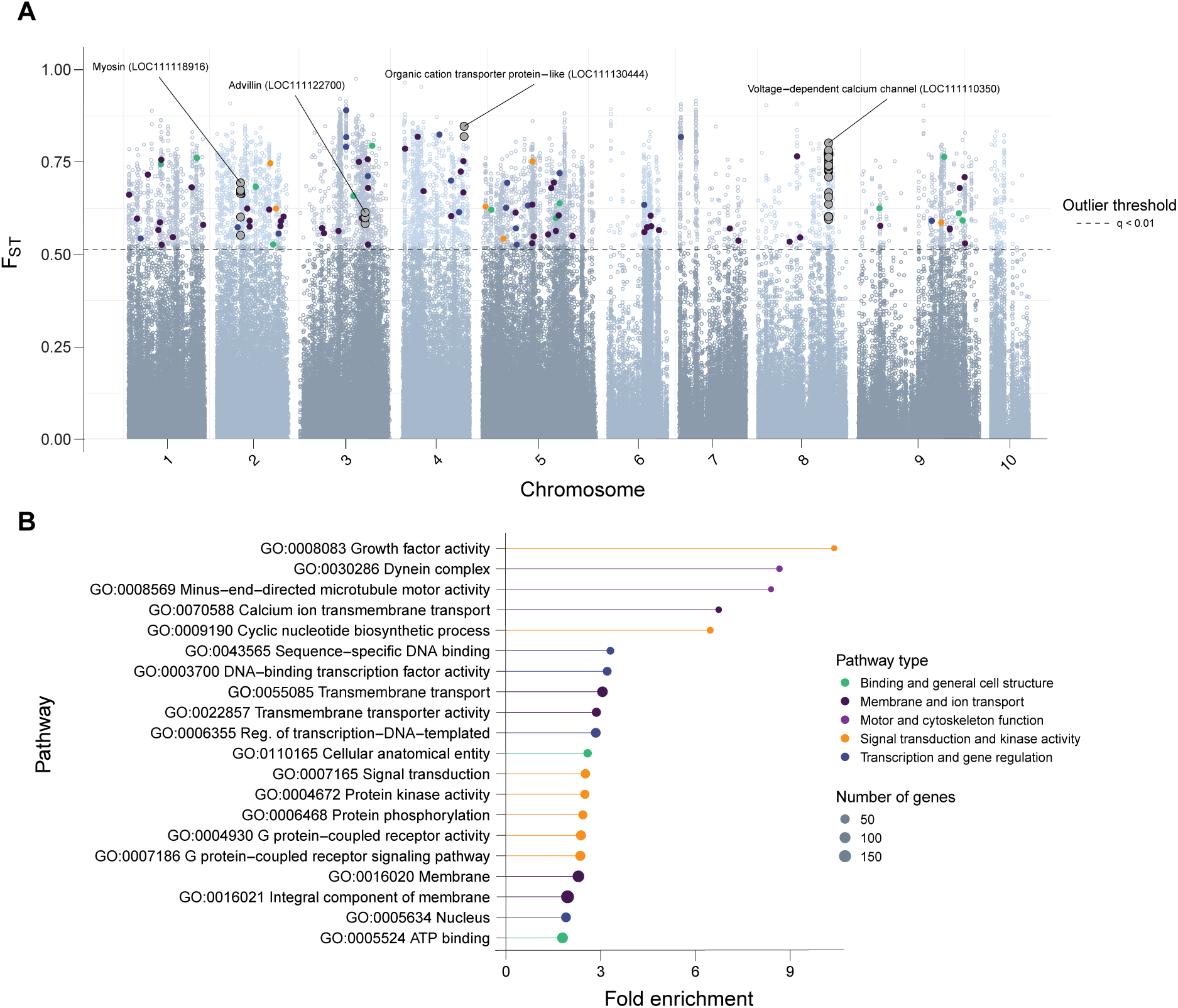
Genome scan. (A) Manhattan plot of *F_ST_* values calculated with OutFLANK on all SNPs. Black dashed line indicates the outlier threshold (q < 0.01). Colored points highlight loci involved in enriched pathways, grouped into five broader categories: binding and general cell structure, membrane and ion transport, motor and cytoskeleton function, signal transduction and kinase activity, and transcription and gene regulation. Grey points highlight loci in genes putatively involved in Dermo resistance (Wang et al. 2025) and osmoregulation. (B) Lollipop plot of fold enrichment values of GO terms corresponding to outlier loci in (A). Point size indicates the number of genes in each pathway. Color corresponds to the same categories as in (A).

The top 20 enriched GO terms involved cell structure, membrane and ion transport, cytoskeletal and motor function, signal transduction, kinase activity, and transcription (Figure 5B). The most enriched pathway was growth factor activity (GO:0008083; fold enrichment = 10.4; 21 genes). Other enriched pathways included calcium ion transmembrane transport (GO:0070588; fold enrichment = 6.7; 36 genes) and transmembrane transporter activity (GO:0022857; fold enrichment = 2.9; 398 genes) (Supp. File 4).

## Discussion

The *local is best, minimize environmental distance*, and *maximize intraspecific diversity* frameworks offer partially conflicting guidelines for transplanting genotypes. Our results provide limited support for any single framework: although a genome scan supported adaptation to disease, heat stress, and salinity, geographically and genetically distant southern groups survived as well as or better than local Chesapeake Bay groups, contradicting *local is best*. Support for *maximize intraspecific diversity* was mixed: polycultures did not outperform monocultures, yet higher parental genetic diversity within monocultures was associated with higher offspring survival (though diversity was confounded with temperature at the EoO). Support for *minimize environmental distance* was also mixed: specific aspects of the EoO (*e.g.,* Salinity_Q10_ and Temperature_Q10_) predicted survival, indicating that historical selection on source populations influenced performance, but relationships between survival and multivariate environmental distance were context-dependent. These patterns were consistent at both field sites despite lower overall survival at the York River.

### Parasite exposure drove site-level differences in survival

The substantially higher mortality at the York River is consistent with the greater disease pressure at that site, where Dermo reached 100% prevalence and MSX reached 53%, compared to 64% Dermo prevalence and no MSX at Lewisetta. Offspring from southern, warmer, and higher-salinity source populations had higher survival at Lewisetta, potentially reflecting historically high Dermo exposure in the Gulf (Ray, 1954). Documented heritable resistance and tolerance to Dermo (Proestou et al., 2019, 2023; Yu & Guo, 2006) could explain the positive relationship between source location salinity and survival at both sites.

Gulf populations showed similar survival to local genotypes in the York River. Because MSX is reported as absent or rare in the Gulf, we would have expected Texas and Louisiana populations to survive less than local genotypes at the York River site. Nevertheless, Gulf populations did show a larger decline in survival between Lewisetta and the York River compared to other populations. Whether southern populations would maintain high survival over longer timescales remains uncertain, as relationships between abiotic factors and disease are complex (Hanley et al. 2019), local adaptation to biotic factors is context-dependent (Briscoe Runquist et al., 2020), and disease-related mortality may manifest seasonally rather than immediately (Ewart & Ford, 1993).

### Evidence for genetic adaptation

Our results suggest genetic adaptation of eastern oysters to varying environmental conditions across their range. Populations differed significantly in survival and growth at common garden sites, but these differences were not explained by large-scale genetic structure. The Gulf and Atlantic formed distinct genetic clusters, yet genome-wide *F_ST_* was low (0.05). Genome scans and gene ontology analyses identified candidate variants potentially underlying these population-level differences.

We detected outlier SNPs in genes previously linked to Dermo resistance (Wang et al., 2025), including *myosin heavy chain*, *advillin*, *heat shock protein 40* (HSP40), and a tumor suppressor. Given that high temperature can intensify Dermo stress (Cook et al., 1998; Hanley et al., 2019; Kachmar et al., 2025), the HSP40 gene (LOC111122700) is particularly notable. Previous work has found upregulation of toll-like receptor and G-protein coupled receptor pathways, suggesting an additional plastic component to immune response (Proestou et al., 2023; Proestou & Sullivan, 2020).

Survival under salinity stress is heritable, with distinct genetic bases for high-and low-salinity tolerance (Allen et al., 2021; McCarty et al., 2020), though populations also buffer environmental stress through physiological plasticity (Eierman and Hare 2016, Sirovy et al. 2023). Standing genetic variation, rather than transgenerational plasticity, appears to be the primary mechanism for persistence under changing conditions (Griffiths et al., 2021). At the genomic level, we found enriched pathways associated with osmoregulation, including transmembrane and calcium ion transport, consistent with prior gene expression results under low salinity stress (Eierman & Hare, 2014; Maynard et al., 2018). Additional enrichment of ion binding, membrane transport, and DNA/RNA replication and repair pathways suggests that salinity adaptation is controlled by many genes of small effect, in accordance with previous QTL data on low salinity survival (McCarty et al., 2022). We note that GO analyses are limited by incomplete annotation and biased toward genes with clear functional hypotheses (Tiffin & Ross-Ibarra, 2014) and an expectation of few large-effect loci when many traits are polygenic (Rausher & Delph, 2015; Rockman, 2012).

### Local is best framework

Local sourcing remains a trusted practice (Broadhurst et al., 2008; O’Brien & Krauss, 2010) due to presumed local adaptation and concerns about outbreeding depression in recipient populations (Frankham et al., 2011). Because “local” is context-dependent, we defined it here as either a wild population from the same estuary and environmental conditions or a selectively bred line that has undergone multiple generations of selection at the common garden site. We found that geographically distant southern populations matched or exceeded local genotypes in survival and growth, while northern populations performed worst. Although we observed GxE interactions across stages (hatchery, nursery, field), we found no evidence for local adaptation, consistent with a previous reciprocal transplant study of Eastern oysters (Hughes et al., 2017). Previous studies in this species have found that southern groups performed better (Burford et al., 2014) or worse than local groups (Proestou et al., 2016; Rawson & Feindel, 2012) when transplanted northward. The high performance of southern genotypes is notable given rapid warming in the Chesapeake Bay (Nardi et al., 2025), which may have already disrupted climate matching at our sites. Our results also provide context for why local genotypes sometimes fail in restoration (Hintenlang et al., 2023; Strickland & Brown, 2024). As climate change continues to disrupt local adaptation (Anderson & Wadgymar, 2020), breeding and restoration programs will increasingly need to consider non-local genotypes.

### Minimize environmental distance framework

Assisted gene flow frameworks emphasize minimizing current or future environmental distance between source and recipient populations to maximize productivity (Aitken & Whitlock, 2013; Supple et al., 2018). We found mixed support for the *minimize environmental distance* framework. On one hand, the relationship between survival and multivariate environmental distance was context-dependent. On the other hand, absolute environmental conditions at the source populations’ environments-of-origin were stronger predictors, and survival was positively associated with temperature and salinity at the source environment. Populations from warmer, saltier environments may have experienced greater historical parasite exposure and thermal stress, and their higher survival may reflect both higher thermal tolerance and disease resistance as discussed above. This suggests that environmental distance is predictive in our system but was not well captured by our multivariate metric. Studies in other systems have found that environment-of-origin is not always a good predictor of performance; for example, environmental data failed to explain yield traits in maize (Li et al., 2025). Future work should incorporate variables such as disease intensity and explore different ways to weight variables in multivariate measures of environmental distance.

### Maximize intraspecific diversity framework

Intraspecific diversity promotes ecosystem function by enhancing resilience, boosting productivity, and stabilizing community structure (Reusch et al. 2005, Crutsinger et al. 2006, Hughes et al. 2008, Abbott et al. 2017, Raffard et al. 2019, 2021). We found mixed support for the *maximize intraspecific diversity* framework. Monocultures with higher heterozygosity and allelic richness had higher survival, but we cannot disentangle the intrinsic benefit of genetic diversity from the source population EoO, as these metrics were confounded with temperature and latitude (i.e., southern populations had higher heterozygosity and allelic richness). Contrary to the framework, polycultures had significantly lower growth and survival than top-performing monocultures.

The two polyculture treatments performed similarly despite their distinct mechanisms for generating diversity. In SEEDMIX (mixed juveniles from non-hybridized monocultures), niche partitioning and differential survival would have largely influenced overall survival (Loreau & Hector, 2001). In HYBRIDMIX (crosses among all source populations), hybrid vigor (Stelkens et al., 2014) and/or outbreeding depression (Frankham et al., 2011; Goto et al., 2011) could have additionally influenced survival. The similar, intermediate performance of these two treatments suggests differential survival was the dominant driver in both. Future parentage analysis of these cohorts could reveal whether the same parents or crosses contributed disproportionately to survival.

Overall, genetic diversity appears to enhance resilience as supported by theory (Fox, 2005; Hughes et al., 2008; Loreau & Hector, 2001), but maladaptive diversity (*e.g.,* the inclusion of poorly performing northern genotypes in our study) can be detrimental. While polycultures are often more productive than monocultures (Huang et al., 2024; Iverson et al., 2014), they may not outperform the most productive monoculture (Morris et al., 2016), a pattern consistent with our results. Previous work in Eastern oysters has shown that the effects of intraspecific diversity are context-dependent: cohorts with higher genetic diversity exhibited improved survival and growth in a hatchery (Hughes et al., 2019), whereas in wild populations, genotypic mixtures enhanced recruitment but sometimes reduced survival (Hanley et al., 2016). Our results suggest that genotypes from a wide geographic range can be combined in polycultures without compromising overall performance, as high-performing genotypes may buffer the effects of weaker or maladapted ones (Brown et al., 2022; Fox, 2005). However, too many disease-susceptible genotypes in a polyculture could drive outbreaks within an otherwise tolerant population.

Balancing the benefits of genetic diversity against the cost of including weaker genotypes remains an important area for future research.

## Conclusion

This study demonstrates that strategies for transplantation cannot be simplified to exclusively environmental or genetic factors, nor is there a clear, successful combination of existing frameworks. Existing recommendations to use regional genotypes to provide an array of traits adaptive in different conditions or seasons in the new environment (a combination of *local is best* and *maximize intraspecific diversity*) (Bucharova et al., 2019; Truskey et al., 2025) may be insufficient to keep pace with rapid environmental change. Assisted gene flow and other strategies for climate change mitigation are being urgently called for as climate change worsens (Nadeau et al. 2024, Aitken et al. 2024, Chakraborty et al. 2024, Baker et al. 2025). Our results show that the Chesapeake Bay will be amenable to assisted gene flow from the south, even from genetically distinct Gulf populations, which may become necessary as estuaries face extreme heat stress (Nardi et al., 2025). While sourcing for aquaculture and restoration remains challenging, we encourage managers to consider a mix of stock sources that are genetically and environmentally equipped for both the pressures of the new environment and for emerging conditions.

## Data and code availability

Data and code are available for peer review in our GitHub repository at https://github.com/DrK-Lo/MVP-H2F-HatcheryField. Upon acceptance of the final manuscript, all data will be archived on BCO-DMO (https://www.bco-dmo.org/project/876610) and code will be archived with Zenodo.

## Author Contributions

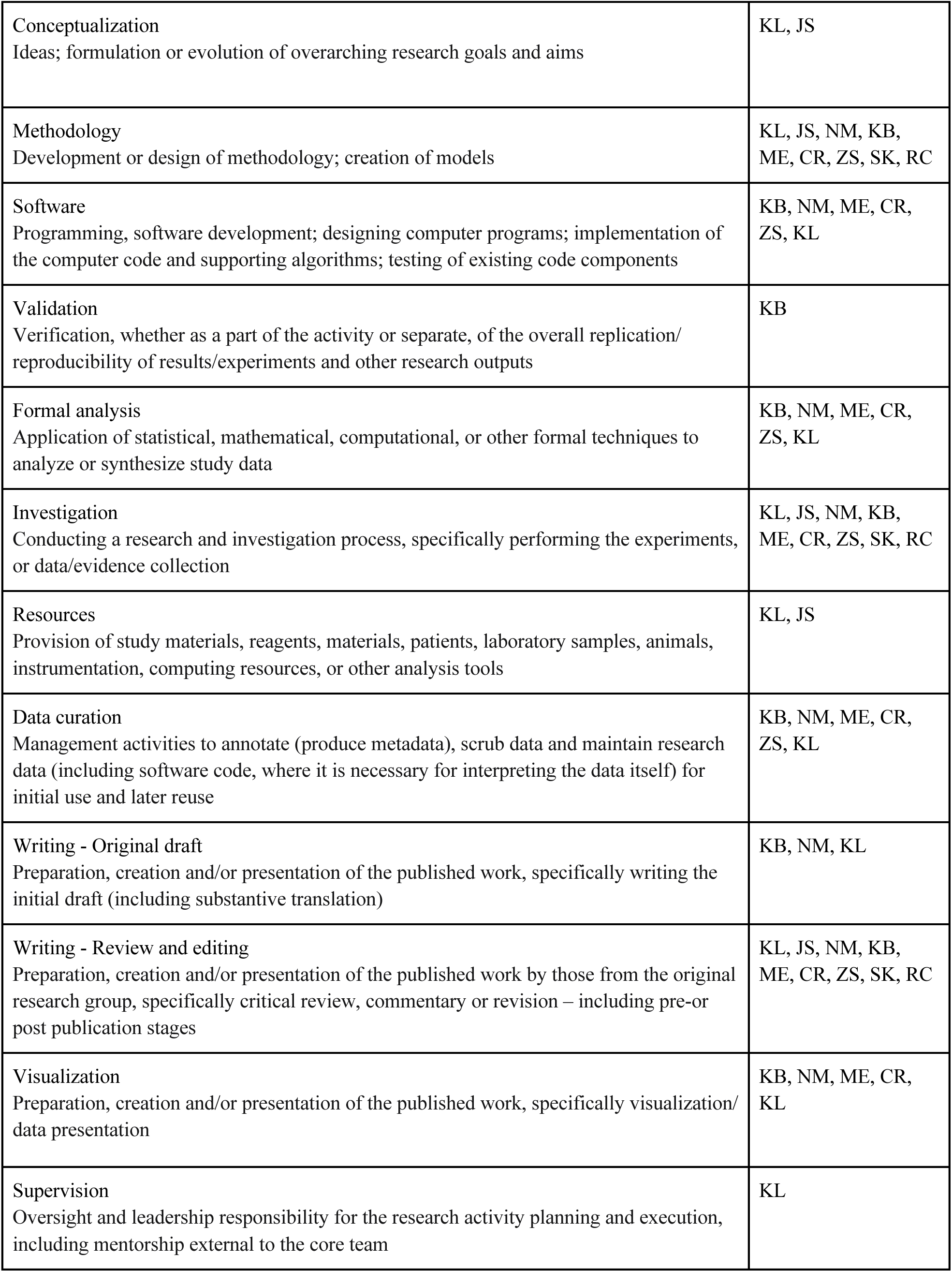
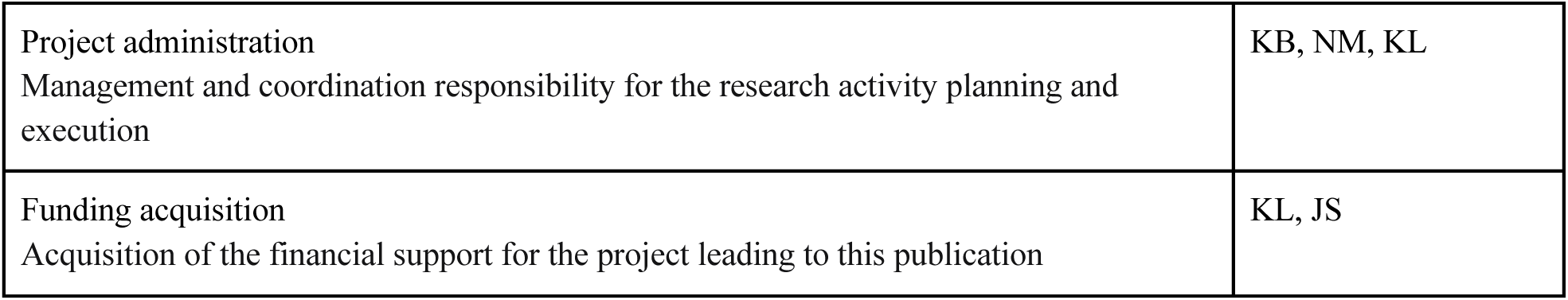

## Funding

This work was funded by a CAREER award from the National Science Foundation to KEL (#2043905).

## Conflict of interest statement

The authors declare no conflicts of interest.

## Supporting information

Supp. File 1 (SNP info)

Supp. File 2 (genotypes)

Supp. File 3 (model results)

Supp. File 4 (GO enrichment)

Supp. File 5 (outlier SNPs)

## Acknowledgements

This work would not have been possible without help from a number of people, to whom we are extremely grateful. We would like to thank the following people who sent us oysters from the source populations: Zachary Olsen, Christine Jensen, Chris Hollenbeck, Willie Cheramie, George Melancon, Hunter Matthews, and Melissa Southworth. We are grateful to the staff at the Aquaculture Technology and Breeding Center for their help conditioning oysters, raising larvae, and monitoring the common gardens: Amanda Chesler-Poole, AJ Verderame, Lauren Gregg, Kate Ritter Sage, Michael Sprague, Haley Uliasz, Meghan Capps, Libby Hoffeditz, and Robin Varney. We would like to thank the following students from Northeastern University for their help in the field and in the lab: Elisabeth Leung, Annie Christie, Annabel Hughes, Lee Fenuccio, Lei Curtis, Zoe Chapman, Anna Eaton, Maya Krattli, Eshna Kulshreshtha, Elizabeth Gouralnik, Fin Li, Nina Johnson, Clara Winguth, Leslie Youtsey, Jack Eynon, Sullivan Newsome, Isabelle Castro, Caela Gilsinian, Quinn Girasek, Julia Grenn, Madison Griffin, and AnnJacob Woodson. Willy Reay provided environmental data for the Lewisetta site. Finally, we are grateful to Jiseon Min, Remy Gatins, and Maddie Armstrong for their helpful comments on the manuscript.

## Notes

https://github.com/DrK-Lo/MVP-H2F-HatcheryField

### Competing Interest Statement

The authors have declared no competing interest.

