## Supplemental information for "Contrasting effects of geographic distance, environmental distance, and intraspecific diversity on the performance of a marine invertebrate in common gardens"

Table of contents


- Supplemental methods 3
  - Broodstock conditioning 3
  - Spawning and fertilization 3
  - Larval rearing 3
  - Survival measurements 4
  - Environmental data 4
    - Environment-of-origin 4
  - DNA extraction 5
  - Genotype filtering 5
  - Statistical analysis 5
- Supplemental files 6
  - Supp file 1: SNP info 6
  - Supp file 2: VCF of the genotype matrix of the full SNP set 6
  - Supp. file 3: All Q2 model results 6
  - Supp file 4: Enriched GO terms from ShinyGO 6
  - Supp. file 5: Full SNP set with outlier status and full annotations 6
- Supplemental tables 7
  - Table S1. Environmental data sources 7
  - Table S2. Parent and common garden environments 10
  - Table S3. ANOVA, survival, final field timepoint 11
  - Table S4. Analysis of deviance, length, final field timepoint 11
  - Table S5. Analysis of deviance, condition index, final field timepoint 11
  - Table S6. Partial Mantel tests 12
- Supplemental figures 14
  - Figure S1. sNMF ancestry of source populations 14
  - Figure S2. Pairwise correlations of environmental quantiles and genetic diversity of source populations 15
  - Figure S3. Shell length by treatment group (end of larval stage - days 15-21, end of nursery stage - day 77) 16
  - Figure S4. Condition index by treatment group (final field timepoint, November 2024) 17
  - Figure S5. Slopes of shell length vs. environmental quantiles, environmental distance, heterozygosity, and allelic richness (all field timepoints) 18
  - Figure S6. Slopes of shell length vs. environmental quantiles, environmental distance,

heterozygosity, and allelic richness (end of larval and nursery stages) 19
Figure S7. Slopes of condition index vs. environmental quantiles, environmental distance, heterozygosity, and allelic richness (all field timepoints) 20
Figure S8. Pairwise correlations of environmental distance and genetic diversity of source populations 21
Figure S9. Survival model residuals vs. environmental distance (final field timepoint, November 2024) 22
Figure S10. Pairwise differences in shell length (final field timepoint, November 2024) 23
References 23

### Supplemental methods

##### *Broodstock conditioning*

Fifty to eighty oysters were collected from each source population in fall 2022 and shipped to the Aquaculture Genetics and Breeding Technology Center (ABC) at the Virginia Institute of Marine Science (VIMS). Non-local animals were held in a quarantine system and fed Shellfish Diet (Reed Mariculture) until conditioning began in February 2023. During conditioning, oysters were held in a flow-through system with 1 µm filtered York River seawater at a slow ramp to 22°C over 4-6 weeks. Oysters were fed live microalgae (*Tetraselmis* and *Pavlova* sp.). Effluent from non-local oysters was treated with 1 µm and 0.5 µm filter bags followed by UV sterilization. Oysters that ripened early were held at 20°C with reduced feeding until spawning.

##### *Spawning and fertilization*

On May 8, 2023, oysters were cleaned, dried, and measured. Twenty parents per population (10 males, 10 females) were strip-spawned following Frank-Lawale et al. (2014) and Proestou et al. (2016). Males were stripped into 50 mL beakers and sperm was brought to volume with seawater and rated for motility and density on a 0-5 scale by a single observer. Females were stripped into 400 mL beakers, eggs were hydrated for several minutes, then rinsed on a 63 µm/20 µm filter stack with 1 µm filtered seawater. Egg density was estimated from 10 µL samples and contributions were equalized to 18 million eggs per group.

For monocultures, pooled eggs were divided among beakers equal to the number of males (typically 10; range 8-10), and each aliquot was fertilized with sperm from a single male. Cell division was confirmed approximately 20 minutes post-fertilization, after which fertilized eggs were recombined and rinsed on a 63 µm/20 µm filter stack to remove excess sperm.

For HYBRIDMIX, eggs from all females across source populations were pooled and divided into one aliquot per experimental male. Each aliquot was fertilized with sperm from a single male, and all fertilized aliquots were then combined. This design maximized the number of unique parental crosses and thus genetic diversity among offspring.

##### *Larval rearing*

Larvae were reared in duplicate tanks (60 L and 200L) per population in aerated, 1 µm filtered seawater at 25-27°C. Cultures were fed live microalgae daily and water was changed every other day. Larval density was reduced at regular intervals by pouring cultures over filter stacks of increasing mesh size. On day 2, larvae from each culture were dropped over a 48 µm/35 µm filter stack which caught healthy D-stage larvae on the top screen and trochophores and undeveloped eggs on the bottom screen. Five larvae were measured to the nearest 5 µm and three replicate counts were taken to estimate survival. Density was adjusted to 10 larvae/mL. This process was repeated on day 7, using a 75 µm filter stack to catch veligers and adjusting to a density of 5 larvae/mL, and on day 14 using a 212 µm filter stack to catch pediveligers/eyed larvae and adjusting to a density of 2 larvae/mL. Excess larvae were either transported (day 2) or shipped overnight on ice (day 14) for preservation in formalin (morphology) and 95% ethanol (genetics).

Timeline of age, life stage, size, and date:

- 0 days; unfertilized egg; 35 µm filter; May 9, 2023
- 2 days; D-stage; 48 µm filter; May 11, 2023
- 8 days; veliger/pediveliger; 75 µm filter; May 17, 2023
- 15-21 days; metamorphosed/eyed; 212 µm filter; May 24-30, 2023

Eyed larvae (>212 µm) were harvested as they developed over days 15-21. Replicate tanks were combined and approximately 150,000 metamorphosed larvae per treatment were set on microcultch in a downwelling system. On day 49, we estimated the proportion of set seed by counting individuals in a known volume and transferred a subset to the nursery setting system. Nursery tanks were filled with raw, unfiltered seawater from the York River. On day 77, we estimated the number of individuals remaining in each population and created SEEDMIX. Oysters were then deployed in the field.

***Survival measurements***

In the hatchery, larval survival was estimated at each density reduction (days 2, 8, 15-21) by counting three replicate samples under a microscope and calculating the mean proportion of live larvae.

Field survival was calculated cumulatively at each assessment (November 2023, May 2024, November 2024). At each timepoint, cumulative survival ($S_t$) for each replicate bag was calculated as:

$$S_t = \frac{n_{live,t}}{n_{live,t} + n_{dead,t}} * S_{t-1} \qquad \text{(Equation 1)}$$

where $n_{live,t}$ is the number of live oysters in a specific bag at timepoint $t$, $n_{dead,t}$ is the number of dead oysters in the bag, and $S_{t-1}$ is the cumulative survival at the previous timepoint. Note that for $t = 1$, $S_{t-1} = 1$. This recursive formula accounts for thinning of the bags as oysters grew by calculating survival relative to oysters present at each timepoint rather than initial stocking density.

***Environmental data***

*Environment-of-origin*

Parental groups were chosen from sites with continuous salinity and temperature monitoring for at least seven months of the year that captured seasonal highs and lows, over a minimum of 8 years, and at an hourly or finer resolution (Table S1). Monitoring buoys were located within one kilometer of each source population, with exceptions noted below.

For W6-ME, the nearest buoy was 3.2 km from the collection site, but since this was a salty estuary without a steep salinity gradient, we used this data for analysis.

For W4-VA and S1-LOLA, high-resolution onsite data were unavailable, so environmental conditions were estimated via interpolation between two data sources and validated against limited onsite datasets: For W4-VA, data came from the VIMS molluscan ecology report (station 435; 37.064°N, 76.563°W) and the NOAA National Data Buoy Center (NDBC) station 44041 (37.211°N, 76.787°W). The VIMS station was geographically closer to the collection site but only recorded data from May through October and failed to capture seasonal variation. The NOAA buoy provided year-round data across multiple years but was located farther from the collection site. VIMS data was recorded once daily; NOAA data were recorded hourly. To integrate these sources, we calculated daily mean temperature and salinity from the hourly NOAA data, then subset both datasets to include only overlapping dates. We calculated the difference in temperature and salinity for each overlapping day, averaged these differences, and applied the resulting values as correction factors to adjust the year-round NOAA data. Corrected values were used in all subsequent environment-of-origin analyses.

For S1-LOLA, data came from a VIMS water quality monitoring station in the Coan River (2021-2025; 37.980°N, 76.462°W) and two NOAA NDBC buoys (2014-2020). Temperature data came from Station LWTV2 (37.995°N, 76.465°W) and salinity data from Station 44042 (38.033°N, 76.335°W), as the nearest buoy recorded only temperature. VIMS data were recorded every 15 minutes; NOAA data were recorded hourly. We calculated daily mean temperature and salinity from both datasets, then followed the same adjustment procedure described above.

##### *DNA extraction*

Samples were randomized on plates, lysed overnight at 56°C, washed, spun, and eluted with 200 µl of low TE buffer (10 mM Tris and 0.1 mM EDTA). Samples were sequenced if they yielded 20 ng/µl of extracted DNA and showed a strong band at > 3k bp on an 2% agarose gel with minimal streaking.

##### *Genotype filtering*

We generated a diploid genotype matrix by coding SNP calls as "0/0" = 0 (homozygous reference), "0/1" or "1/0" = 1 (heterozygous), or "1/1" = 2 (homozygous alternate) using `vcfR` v1.15.0. The genotype matrix was initially filtered to retain nuclear SNPs and remove SNPs with > 10% missing data across all loci and all individuals. SNPs were then filtered for minor allele frequency (MAF) > 0.05.

##### *Statistical analysis*

For general linear models and general linear mixed models, we assessed model assumptions of normality and homoscedasticity using Q-Q plots and scale-location plots, respectively. For multiple regression models, we assessed multicollinearity and removed a collinear term when the pairwise Pearson's correlation (r) exceeded 0.7. In the case of multiple tests, p-values were adjusted for multiple comparisons using the Benjamini and Hochberg method (Benjamini and Hochberg 1995).

#### Supplemental files

See README for detail on each file: https://github.com/DrK-Lo/MVP-H2F-HatcheryField/tree/main/supp_files

Note to reviewers: all supplemental files will be archived on our project page on BCO-DMO at https://www.bco-dmo.org/project/876610

Supp. file 1: SNP info
Name, sequence, position etc. for SNPs that passed missingness and MAF (full SNP set). Also includes a column that indicates if the SNP is in the thinned SNP set.
Supp. file 2: VCF of the genotype matrix of the full SNP set
Supp. file 3: All Q2 model results
Supp. file 4: Enriched GO terms from ShinyGO
Supp. file 5: Full SNP set with outlier status and full annotations

#### Supplemental tables

##### ***Table S1. Environmental data sources***

Table S1. Location, data access information, and resolution of environmental data sources for each environment-of-origin and common garden site.

| **Environment type** | **Site name** | **Latitude** | **Longitude** | **Data origin** | **Data access** | **Span of data (years)** | **Months of data per year** | **Data interval** |
|---|---|---|---|---|---|---|---|---|
| Environment-of-origin (EoO) | W1-TX | 28.084 | -97.201 | NERR Centralized Data. Mission Aransas - Copano Bay West MARCWWQ | https://cdmo.baruch.sc.edu//dges/ | 2007 - 2021 | 12 | 15 minutes |
| EoO | W2-LA | 29.249 | -90.921 | USGS Water Data Sister Lake - 07381349 | https://waterdata.usgs.gov/monitoring-location/07381349/#parameterCode=00065&period=P7D | 2007 - 2022 | 12 | 15 minutes, 30 minutes, or 1 hour |
| EoO | W3-FL | 30.441 | -81.439 | National Parks Service Continuous Water Data - Timucuan Preserve | https://irma.nps.gov/AQWebPortal/Data/Location/Summary/Location/TIMUking01/Interval/Latest | 2005 - 2022 | 12 | 30 minutes |
| EoO | W4-VA | 37.211 | -76.787 | NOAA National Data Buoy Center (NDBC), Chesapeake Bay Interpretive Buoy System - 44041 | https://www.ndbc.noaa.gov/station_page.php?station=44041 | 2008 - 2019 | 12 | 1 hour |
| | | 37.064 | -76.563 | VIMS - Molluscan ecology report, provided by Melissa Southworth | https://www.vims.edu/research/units/labgroups/molluscan_ecology/publications/ | 2010 - 2023 | 4 | 7 days or 14 days |
| | | Compilation: NDBC values corrected according to data from VIMS. Compiled data range: 2012 - 2019. | | | | | | |

| | | | | | | | | |
|---|---|---|---|---|---|---|---|---|
| EoO | S1-LOLA | 37.980 | -76.462 | VIMS, provided by William Reay | Private access | 2023 - 2024 | 12 | 15 minutes |
| | | 38.033 | -76.335 | NDBC, Chesapeake Bay Interpretive Buoy System - 44042 | https://www.ndbc.noaa.gov/station_page.php?station=44042 | 2014 - 2024 | 12 | 1 hour |
| | | 37.995 | -76.465 | NDBC, National Ocean Service - LWTV2 - 8635750 | https://www.ndbc.noaa.gov/station_page.php?station=lwtv2 | 2014 - 2024 | 2 | 6 minutes |
| | | Compilation: VIMS salinity data corrected according to NDBC station 44042, VIMS temperature data corrected according to NDBC station 8635750. Compiled data range: 2014 - 2024. | | | | | | |
| EoO | S2-DEBY | 37.247 | -76.499 | Virginia Estuarine and Coastal Observing System, VIMS | https://vecos.vims.edu/dashboard?station=YRK005.40 | 2003 - 2024 | 12 | 15 minutes |
| EoO | W5-NH | 43.052 | -70.912 | NERR Centralized Data. Great Bay - Squamscott River GRBSQWQ | https://cdmo.baruch.sc.edu//dges/ | 1997 - 2022 | 7, 8, 9 or 12 | 15 minutes or 30 minutes |
| EoO | W6-ME | 43.986 | -69.55 | University of Maine | http://maine.loboviz.com/ | 2015 - 2023 | 7, 8, or 9 | 1 hour |
| Common garden | Lewisetta | 37.980 | -76.462 | VIMS, provided by William Reay | Private access | 2023 - 2024 | 12 | 15 minutes |
| Common garden | York River | 37.247 | -76.499 | Virginia Estuarine and Coastal Observing System, VIMS | https://vecos.vims.edu/dashboard?station=YRK005.40 | 2023 - 2024 | 12 | 15 minutes |

***Table S2. Parent and common garden environments***

Table S2. Mean annual, 0.1 quantile (Q10), and 0.9 quantile (Q90) values for salinity (ppt) and temperature (°C) at each environment-of-origin and common garden site.

| **Site name** | **Mean annual salinity (ppt)** | **$Salinity_{Q10}$** | **$Salinity_{Q90}$** | **Mean annual temperature (°C)** | **$Temperature_{Q10}$** | **$Temperature_{Q90}$** |
|---|---|---|---|---|---|---|
| W1-TX | 20.9 | 5.9 | 36.7 | 23.1 | 13.8 | 30.5 |
| W2-LA | 10.5 | 3.0 | 18 | 22.9 | 13.2 | 30.7 |
| W3-FL | 32.4 | 27.9 | 36 | 22.2 | 14.6 | 29.3 |
| W4-VA | 15.2 | 12.2 | 19.1 | 18.5 | 6.1 | 28.1 |
| S1-LOLA | 12.7 | 8.7 | 16.6 | 15.9 | 5.0 | 27.7 |
| S2-DEBY | 19.3 | 15.8 | 22.4 | 17.1 | 6.5 | 27.5 |
| W5-NH | 18.6 | 5.9 | 28.6 | 16.7 | 7.2 | 24.2 |
| W6-ME | 29.5 | 26.3 | 31.5 | 15.3 | 7.6 | 21.5 |
| Lewisetta | 13.1 | 8.3 | 16.5 | 18.4 | 7.2 | 28.2 |
| York | 19.8 | 15.4 | 22.9 | 18.1 | 8 | 27.4 |

*Table S3. ANOVA, survival, final field timepoint*

Table S3. Results of a two-way ANOVA examining the effects of common garden site, treatment group, and their interaction on survival at the final field timepoint.

| **Variable** | **Type 3 SS** | **df** | **F** | **p** |
|---|---|---|---|---|
| Common garden site | 0.10500 | 1 | 26.950 | < 0.001 |
| Treatment group | 1.71580 | 9 | 48.930 | < 0.001 |
| Common garden site:Treatment group | 0.39027 | 9 | 11.129 | <0.001 |

*Table S4. Analysis of deviance, length, final field timepoint*

Table S4. Results of an analysis of deviance examining the effects of common garden site, treatment group, and their interaction (with bag as a random effect) on length at the final field timepoint.

| **Explanatory variable** | $\chi^2$ | **df** | **p** |
|---|---|---|---|
| Common garden site | 8.6348 | 1 | 0.003 |
| Treatment group | 238.4768 | 9 | < 0.001 |
| Common garden site:Treatment group | 73.2551 | 9 | < 0.001 |

*Table S5. Analysis of deviance, condition index, final field timepoint*

Table S5. Results of an analysis of deviance examining the effects of common garden site, treatment group, and their interaction (with bag as a random effect) on condition index at the final field timepoint.

| **Variable** | $\chi^2$ | **df** | **p** |
|---|---|---|---|
| Common garden site | 15.934 | 1 | < 0.001 |
| Treatment group | 22.097 | 9 | 0.00857 |
| Common garden site:Treatment group | 15.856 | 9 | 0.070 |

#### *Table S6. Partial Mantel tests*

Table S6. Results of partial Mantel tests examining the effect of either genetic distance ($F_{ST}$) while controlling for environmental distance or the effect of environmental distance while controlling for genetic distance on all response variables used in this study: survival and length in the hatchery, in the nursery, and at all field timepoints.

| Explanatory variable | Variable partialed out | Response variable | p-value | Adjusted p-value |
|---|---|---|---|---|
| Genetic distance ($F_{st}$) | Environmental distance | Day 21 length | 0.555 | 0.724 |
| | | Day 77 length | 0.297 | 0.558 |
| | | $t_1$ Lewisetta length | 0.0307 | 0.123 |
| | | $t_1$ Lewisetta survival | 0.0189 | 0.113 |
| | | $t_1$ York length | 0.767 | 0.834 |
| | | $t_1$ York survival | 0.142 | 0.377 |
| | | $t_2$ Lewisetta length | 0.00780 | 0.0585 |
| | | $t_2$ Lewisetta survival | 0.00720 | 0.0585 |
| | | $t_2$ York length | 0.675 | 0.788 |
| | | $t_2$ York survival | 0.202 | 0.405 |
| | | $t_3$ Lewisetta length | 0.0226 | 0.113 |
| | | $t_3$ Lewisetta survival | 0.0329 | 0.123 |
| | | $t_3$ York length | 0.361 | 0.636 |
| | | $t_3$ York survival | 0.199 | 0.405 |
| Environmental distance | Genetic distance ($F_{st}$) | Day 21 length | 0.196 | 0.405 |
| | | Day 77 length | 0.806 | 0.834 |
| | | $t_1$ Lewisetta length | 0.789 | 0.834 |

| | | | | |
|---|---|---|---|---|
| | | $t_1$ Lewisetta survival | 0.481 | 0.688 |
| | | $t_1$ York length | 0.447 | 0.671 |
| | | $t_1$ York survival | 0.151 | 0.377 |
| | | $t_2$ Lewisetta length | 0.921 | 0.921 |
| | | $t_2$ Lewisetta survival | 0.412 | 0.671 |
| | | $t_2$ York length | 0.593 | 0.741 |
| | | $t_2$ York survival | 0.116 | 0.377 |
| | | $t_3$ Lewisetta length | 0.533 | 0.724 |
| | | $t_3$ Lewisetta survival | 0.425 | 0.671 |
| | | $t_3$ York length | 0.683 | 0.788 |
| | | $t_3$ York survival | 0.144 | 0.377 |

Supplemental figures

***Figure S1. sNMF ancestry of source populations***

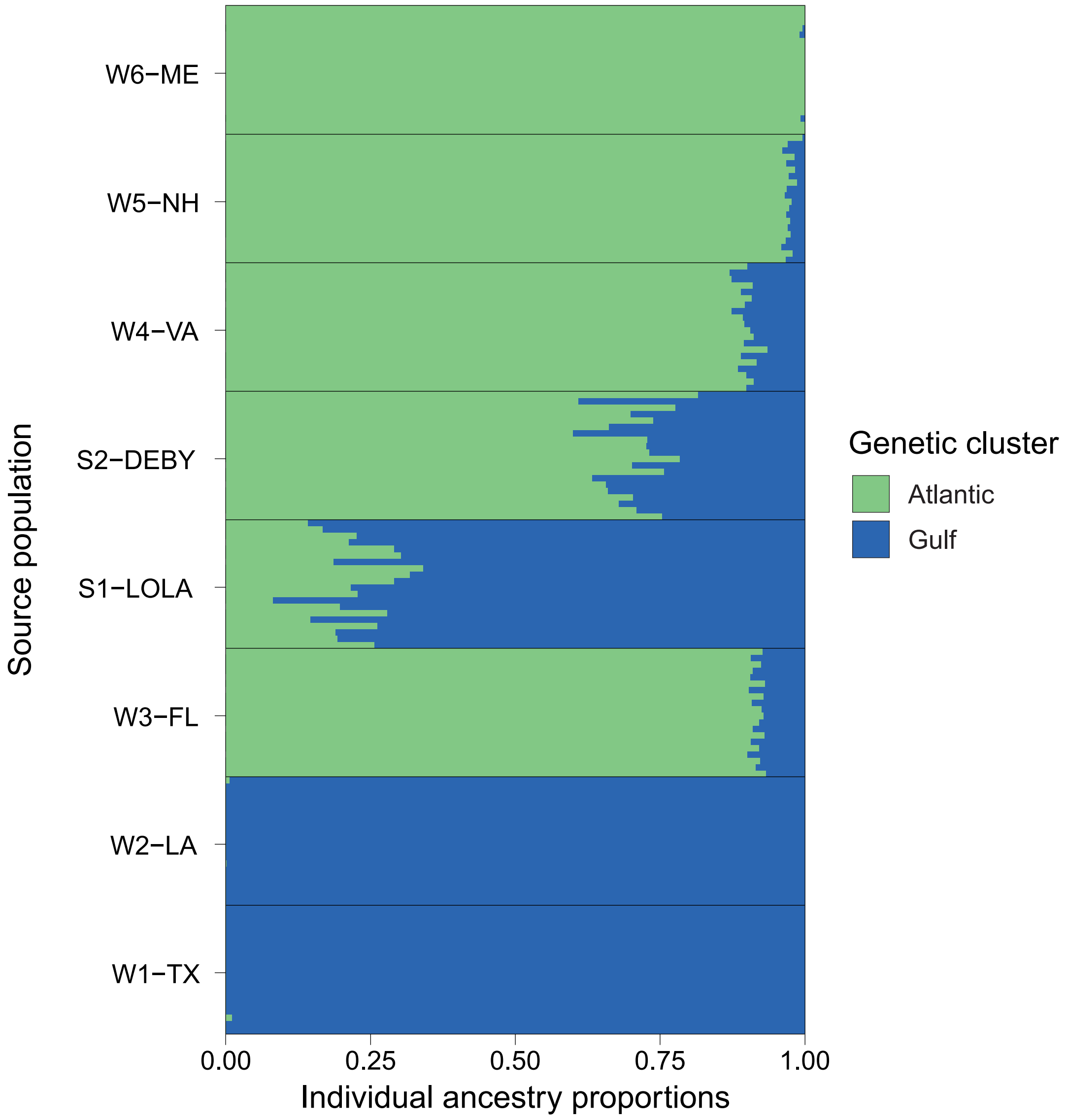


Figure S1. sNMF ancestry analysis performed on SNPs thinned for LD with K = 2 and 10 repetitions. Each row represents one individual; green bars indicate Atlantic ancestry and blue bars indicate Gulf ancestry.

***Figure S2. Pairwise correlations of environmental quantiles and genetic diversity of source populations***

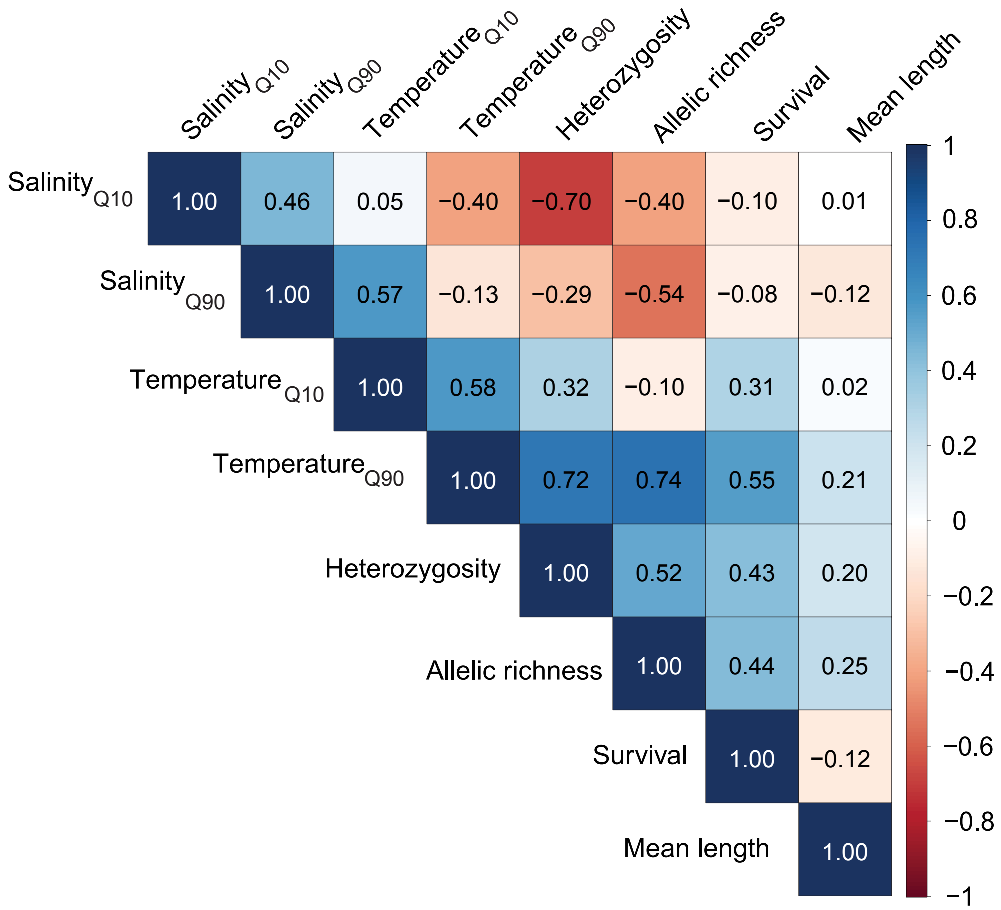


Figure S2. Pairwise correlations among response variables (survival and mean length) and explanatory variables (0.1 and 0.9 quantiles of salinity and temperature at the source populations' environment-of-origin, heterozygosity of source populations, and allelic richness of source populations) at each common garden site: (A) Lewisetta, and (B) the York River.

***Figure S3. Shell length by treatment group (end of larval stage - days 15-21, end of nursery stage - day 77)***

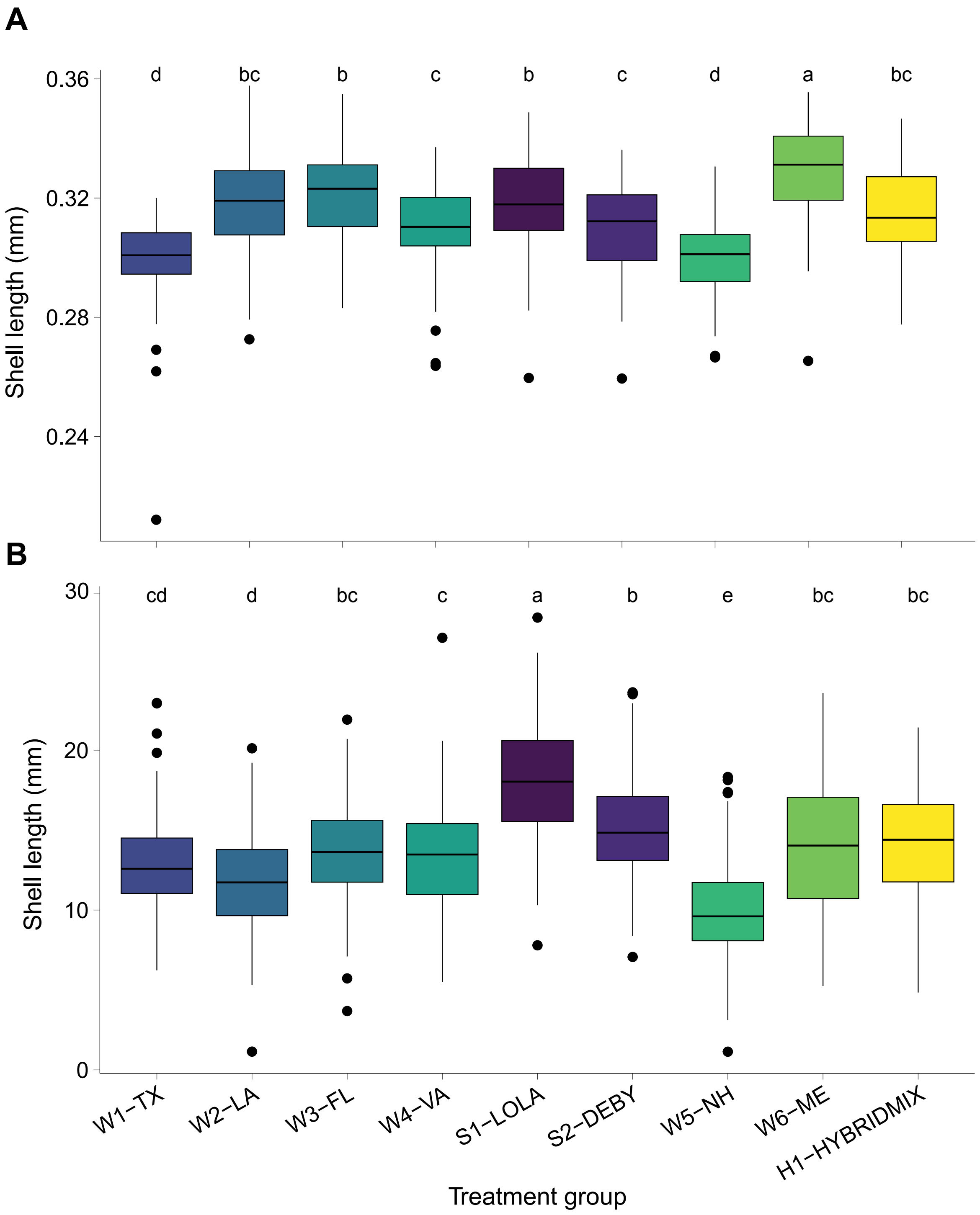


Figure S3. Boxplot of shell length (mm) of oysters on (A) days 15-21, the end of the larval stage, and (B) day 77, the final day of nursery culture before deployment in the field. Letters indicate significant ($p < 0.05$) pairwise differences between treatment groups.

***Figure S4. Condition index by treatment group (final field timepoint, November 2024)***

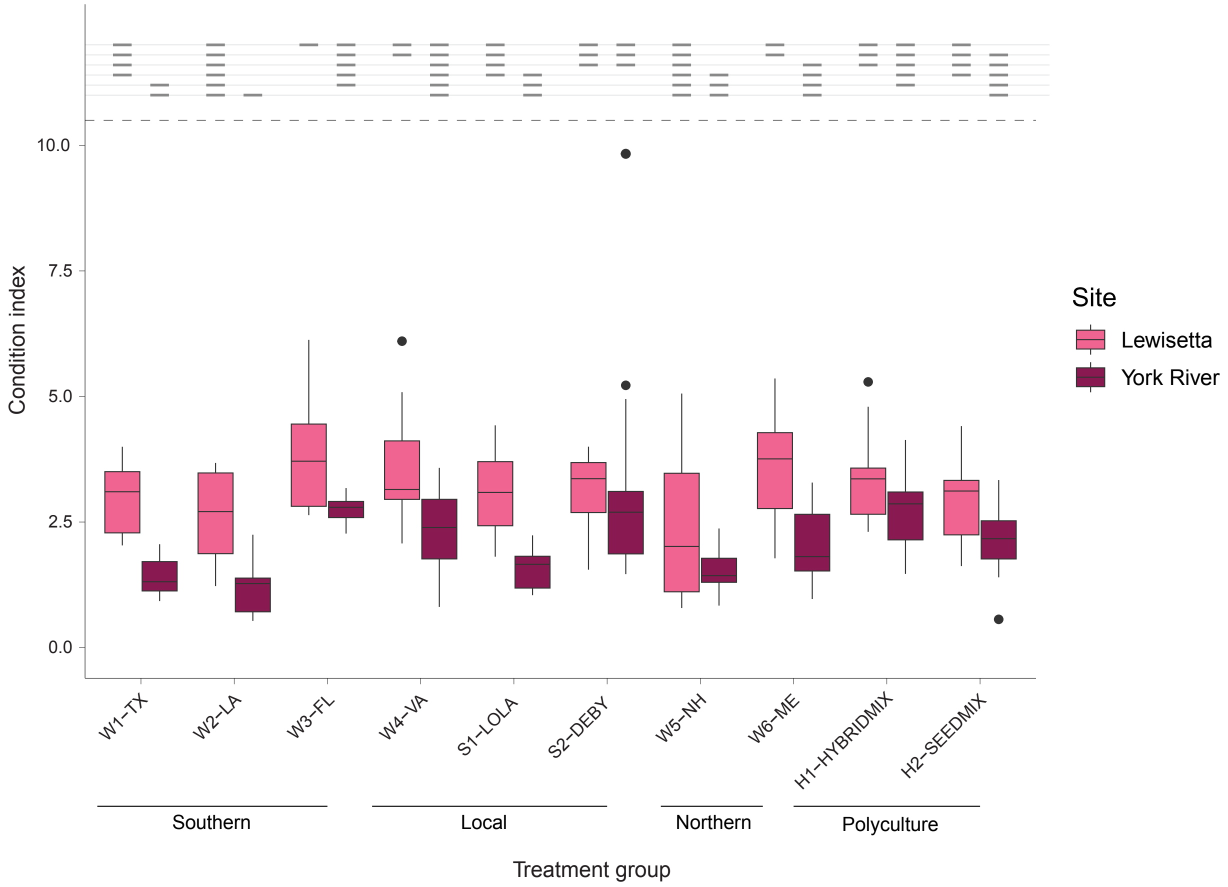


Figure S4: Boxplot of condition index of each treatment group at the final field timepoint (November 2024) at the Lewisetta (lighter boxes) and York River (dark boxes) common garden sites. Horizontal lines above the boxes indicate significant pairwise differences: groups that share a line at the same height do not have a statistically significant difference.

#### *Figure S5. Slopes of shell length vs. environmental quantiles, environmental distance, heterozygosity, and allelic richness (all field timepoints)*

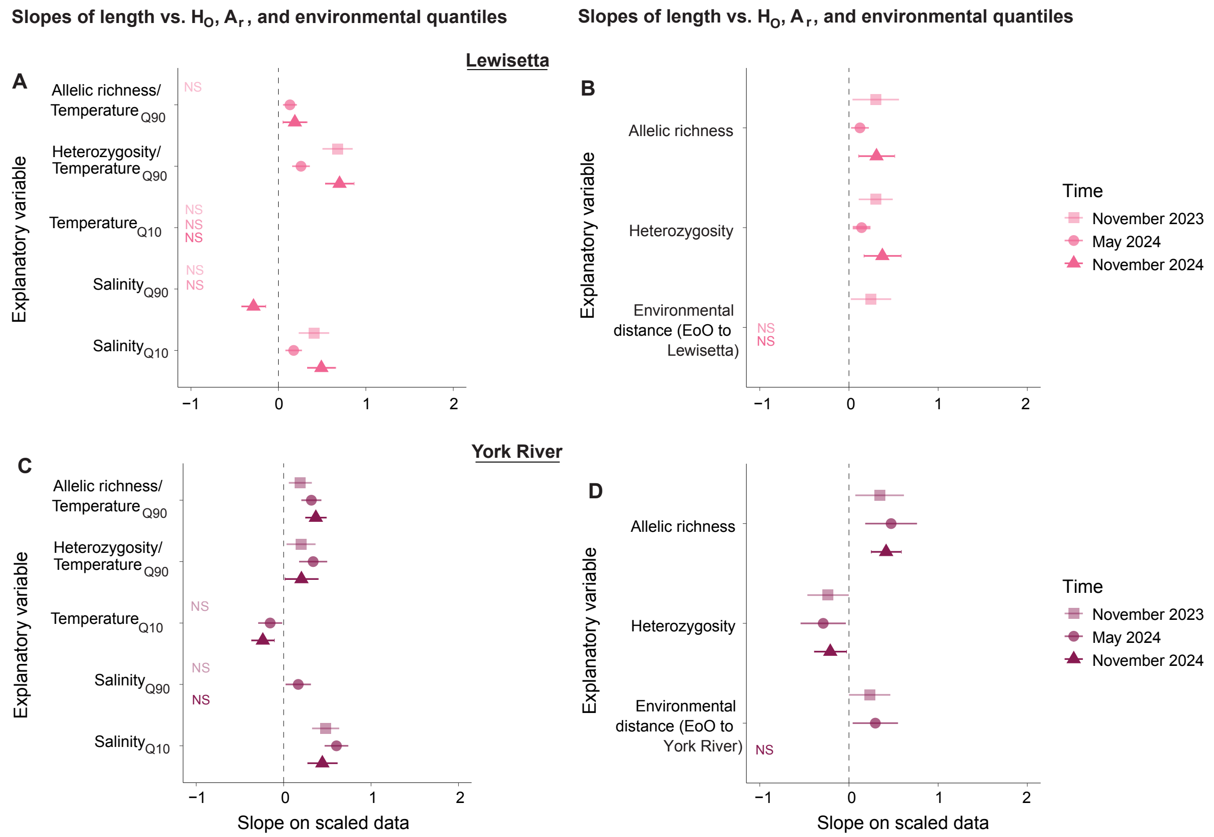


Figure S5. Slope ± 2SE of scaled length at Lewisetta at three field timepoints vs. (A) genetic diversity variables and absolute conditions at source populations' environments-of-origin (EoO), and vs. (B) scaled genetic diversity variables and environmental distance from EoOs to the Lewisetta common garden site. Slope ± 2SE of scaled length at the York River at three field timepoints vs. (C) genetic diversity variables and absolute conditions at source populations' EoO, and vs. (D) scaled genetic diversity variables and environmental distance from EoOs to the York River common garden site.

***Figure S6. Slopes of shell length vs. environmental quantiles, environmental distance, heterozygosity, and allelic richness (end of larval and nursery stages)***

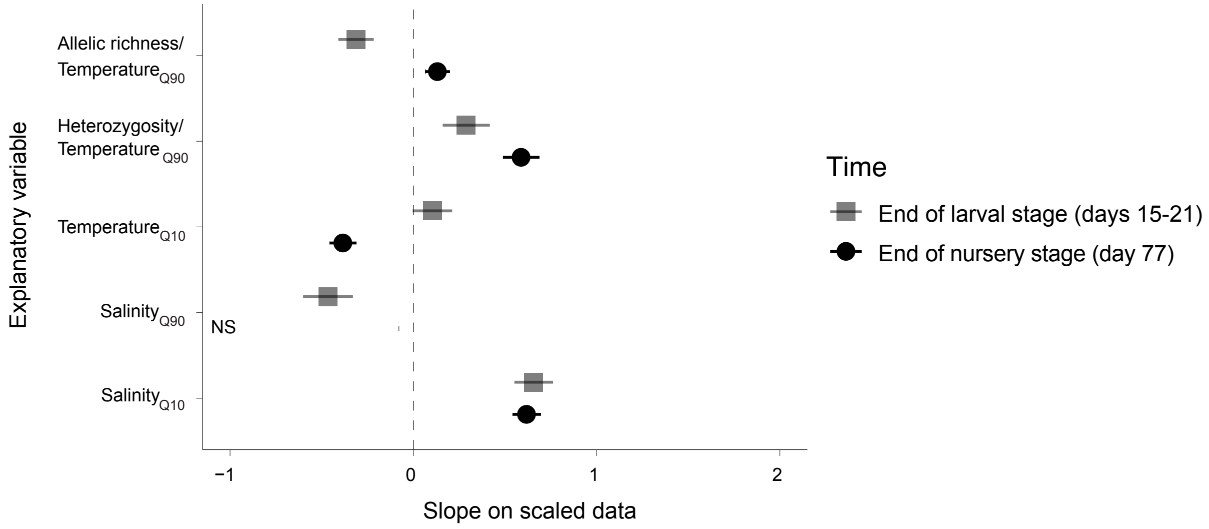


Figure S6. Slope ± 2SE of scaled length at the end of the larval stage (days 15-21) and the end of the nursery stage (day 77) vs. genetic diversity variables and absolute conditions at source populations' environments-of-origin (EoO).

#### *Figure S7. Slopes of condition index vs. environmental quantiles, environmental distance, heterozygosity, and allelic richness (all field timepoints)*

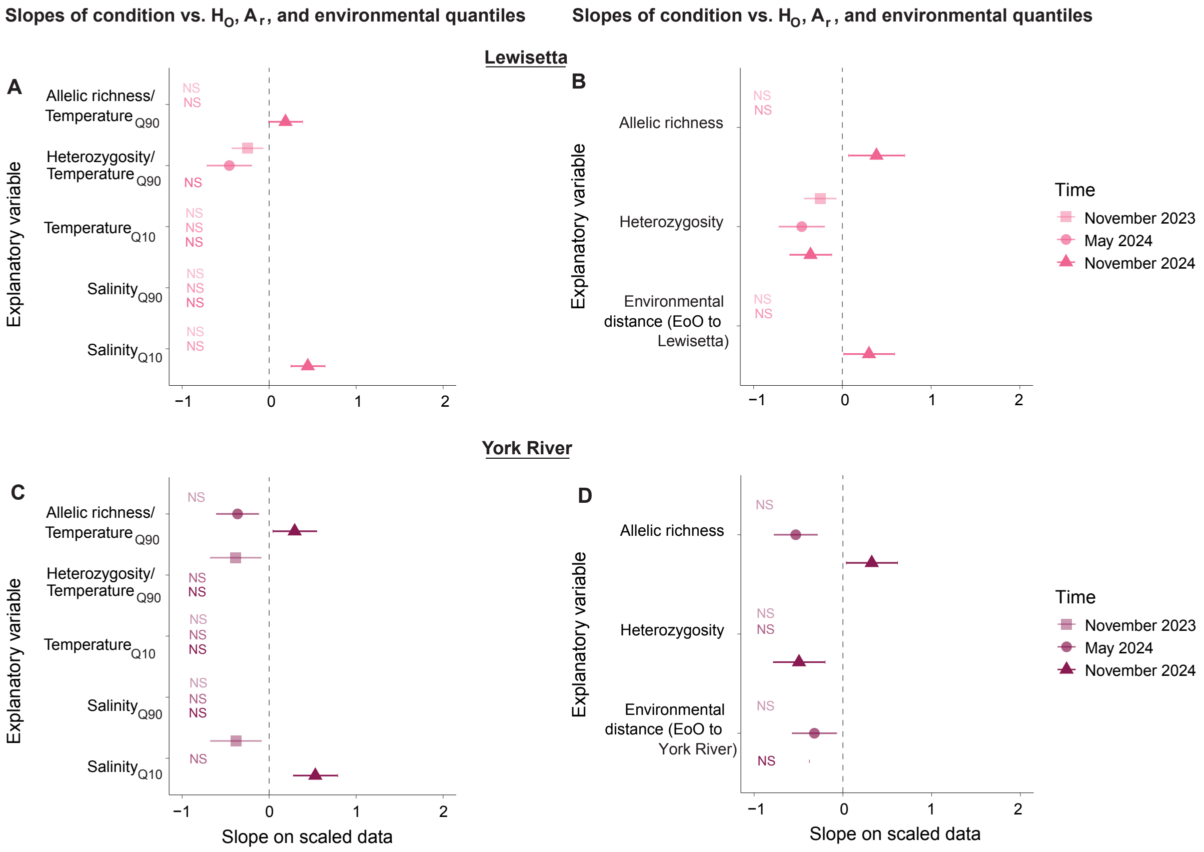


Figure S7. Slope ± 2SE of scaled condition index at Lewisetta at three field timepoints vs. (A) genetic diversity variables and absolute conditions at source populations' environments-of-origin (EoO), and vs. (B) scaled genetic diversity variables and environmental distance from EoOs to the Lewisetta common garden site. Slope ± 2SE of scaled condition index at the York River at three field timepoints vs. (C) genetic diversity variables and absolute conditions at source populations' EoO, and vs. (D) scaled genetic diversity variables and environmental distance from EoOs to the York River common garden site.

***Figure S8. Pairwise correlations of environmental distance and genetic diversity of source populations***

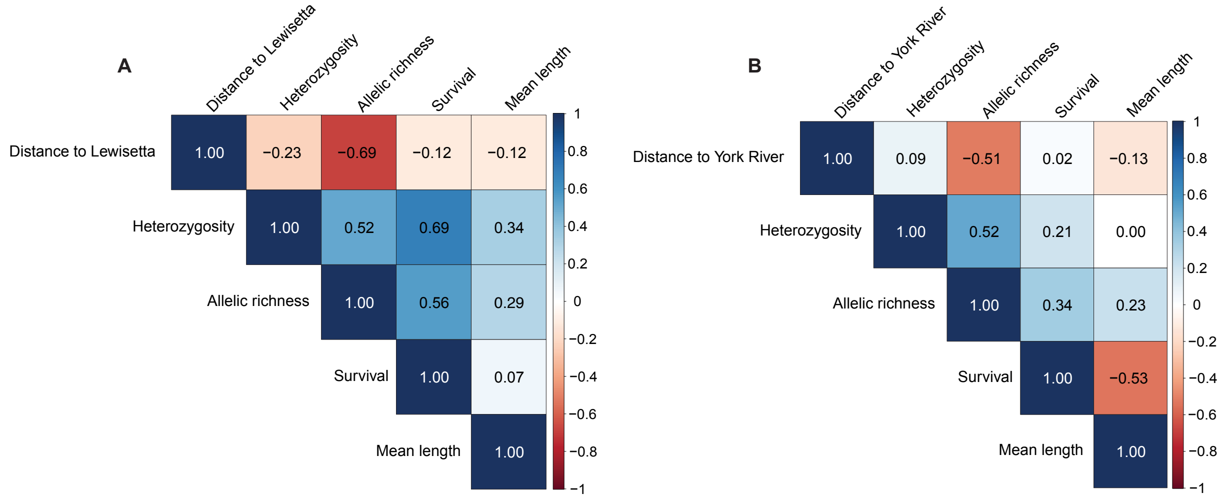


Figure S8. Pairwise correlations among response variables (survival and mean length) and explanatory variables (environmental distance between common garden site and the source populations' environment-of-origin, heterozygosity of source populations, and allelic richness of source populations) at each common garden site: (A) Lewisetta, and (B) the York River.

***Figure S9. Survival model residuals vs. environmental distance (final field timepoint, November 2024)***

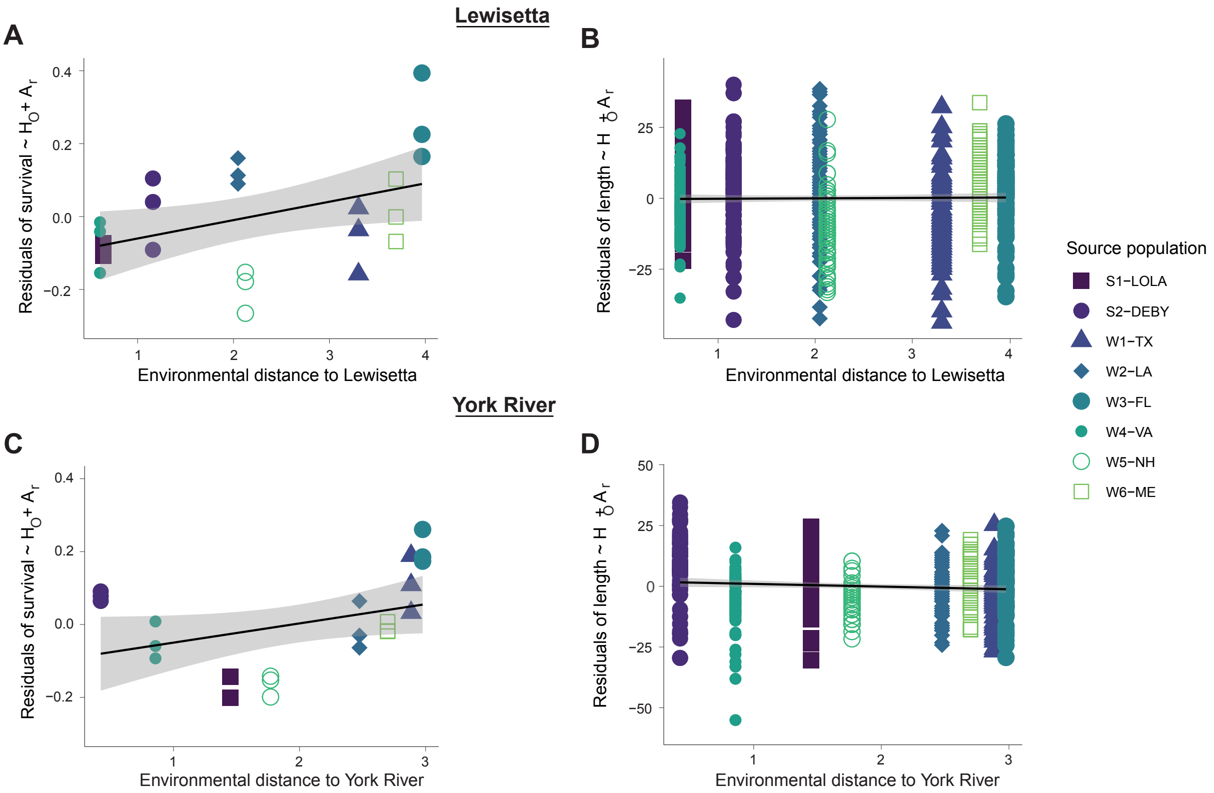


Figure S9. Residuals of linear models relating (A) survival or (B) shell length at the final field timepoint to heterozygosity and allelic richness vs. environmental distance to Lewisetta. Residuals of linear models relating (C) survival or (D) shell length at the final field timepoint to heterozygosity and allelic richness vs. environmental distance to the York River.

***Figure S10. Pairwise differences in shell length (final field timepoint, November 2024)***

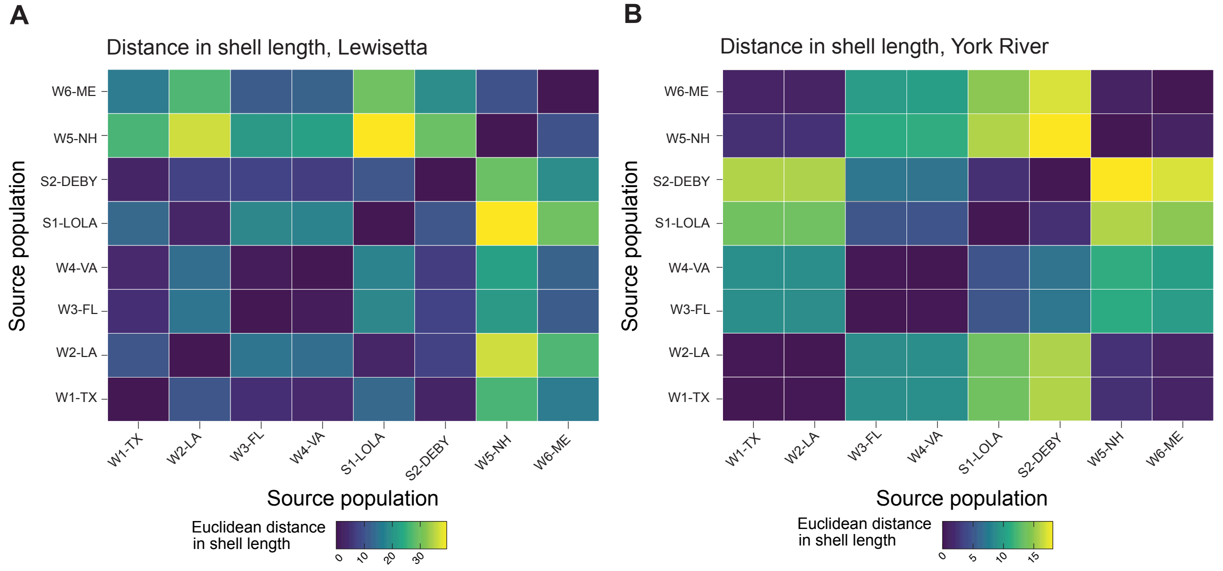


Figure S10. Euclidean distance in shell length between treatment groups at the final field timepoint at (A) Lewisetta and (B) the York River. Darker colors indicate small pairwise differences and lighter colors indicate greater pairwise differences.
